# The somatic genetic and epigenetic mutation rate in a wild long-lived perennial *Populus trichocarpa*

**DOI:** 10.1101/862623

**Authors:** Brigitte T. Hofmeister, Johanna Denkena, Maria Colomé-Tatché, Yadollah Shahryary, Rashmi Hazarika, Jane Grimwood, Sujan Mamidi, Jerry Jenkins, Paul P. Grabowski, Avinash Sreedasyam, Shengqiang Shu, Kerrie Barry, Kathleen Lail, Catherine Adam, Anna Lipzen, Rotem Sorek, Dave Kudrna, Jayson Talag, Rod Wing, David W. Hall, Gerald A. Tuskan, Jeremy Schmutz, Frank Johannes, Robert J. Schmitz

## Abstract

**Background:** Plants can transmit somatic mutations and epimutations to offspring, which in turn can affect fitness. Knowledge of the rate at which these variations arise is necessary to understand how plant development contributes to local adaption in an eco-evolutionary context, particularly in long-lived perennials.

**Results:** Here, we generated a new high-quality reference genome from the oldest branch of a wild *Populus trichocarpa* tree with two dominant stems which have been evolving independently for 330 years. By sampling multiple, age-estimated branches of this tree, we used a multi-omics approach to quantify age-related somatic changes at the genetic, epigenetic and transcriptional level. We show that the per-year somatic mutation and epimutation rates are lower than in annuals and that transcriptional variation is mainly independent of age divergence and cytosine methylation. Furthermore, a detailed analysis of the somatic epimutation spectrum indicates that transgenerationally heritable epimutations originate mainly from DNA methylation maintenance errors during mitotic rather than during meiotic cell divisions.

**Conclusion:** Taken together, our study provides unprecedented insights into the origin of nucleotide and functional variation in a long-lived perennial plant.

## BACKGROUND

The significance of somatic mutations, i.e., variations in DNA sequence that occur after fertilization, in long-lived plant and animal species have been a point of debate and investigation for the past 30 years [1–4]. It has been hypothesized that the evolutionary consequences of such mutations are likely even more profound in woody perennial plants, where undifferentiated meristematic cells produce all above-ground and below-ground structures. As meristems undergo constant cell division throughout the lifetime of a plant, somatic mutations arising in meristems may result in genetic differences being passed onto progeny cells [5–8]. The accumulation of somatic mutations can thus lead to genetic and occasionally also phenotypic divergence among vegetative lineages within the same individual. In trees, for instance, different branches have been shown to differ in their responses to pest and pathogen attack, alternate reactions to drought and/or nutrient availability, or dissimilar demands for photosynthate material, even within the same individual [9]. Beyond the impact of point mutations and small insertions/deletions on gene function, alterations in chromatin structure and DNA methylation might also impact gene expression variation.

Phenotypic variation has been attributed to somatic mutations in several perennial plants, including the derivation of Nectarines in peach [10] and the origin of modern grape cultivars (*Vitis vinifera* L.) [11]. In *Populus tremuloides,* somatic mutations have been hypothesized as the cause for variation in DNA markers among individual ramets of a single genotype [12]. Initial attempts to demonstrate within-tree mosaicism using genetic markers [13], showed at low-resolution that the degree of intra-tree variability was positively correlated with the physical distance between sampled branches. More recently, work in oak (*Quercus rubur*) has documented variation in DNA sequence among an independent sampling of alternate branches from a single genotype [14, 15]. They estimated a fixed mutation rate of 4.2 - 5.2 × 10^−8^ substitutions per locus per generation, which is only within one order of magnitude of the rate observed in the herbaceous annual plant *Arabidopsis thaliana* [16]. These results are consistent with an emerging hypothesis that the per-unit-time mutation rate of perennials is much lower than in annuals to delay mutational meltdown [17, 18] and this lower rate is accomplished by limiting the number of cell divisions between the meristem and the new branch [19]. Additional recent studies have also revealed similar rates of spontaneous mutations in a range of species including perennials [18]. Regardless of the rate of mutation, the frequency of deleterious mutations in woody plants is high, which is hypothesized to reduce survival of progeny resulting from inbreeding and favor outcrossing as is observed in many forest trees [20, 21].

Similar to genetic mutations, phenotypic variation can be caused by epigenetic variation such as stable changes in cytosine methylation or epimutations [22]. Cytosine methylation is a covalent base modification that is inherited through both mitotic and meiotic cell divisions in plants [23]. It occurs in three sequence contexts, CG, CHG, and CHH (H = A, T, or C) and the pattern and distribution of methylation at these different contexts is predictive of its function in genome regulation [24]. Spontaneous changes in methylation independent of genetic changes can lead to phenotypic changes [25]. Well-characterized examples in plants include the peloric phenotype in toadflax (*Linaria vulgaris*), the colorless non-ripening phenotype in tomato (*Solanum lycopersicum*), and the mantled phenotype in oil palm (*Elaeis guineensis*) [26–28].

Once established, epimutations can stably persist or be inherited across generations. For example, the reversion rate from the colorless non-ripening epimutant allele to wild type is about 1 in 1000 per generation in tomato [27]. Studies in *A. thaliana* mutation accumulation lines have documented that the vast majority (91-99.998%) of methylated regions in the genome are stably inherited across generations; only a small subset of the methylome shows variation among mutation accumulation lines [29–31]. Estimates in *A. thaliana* indicate that the spontaneous methylation gain and loss rates at CG sites are 2.56 × 10^−4^ and 6.30 × 10^−4^ per generation per haploid methylome, respectively [32]. Despite the wealth of knowledge about transgenerational methylation inheritance, very little is known about somatic epimutations, especially in long-lived perennial species. Previous studies have been limited by resolution and time. Heer *et al.* observed no global methylation changes and no consistent variation in gene body methylation associated with growth conditions of Norway spruce [33]. Several studies have linked stress conditions to differential methylation in perennials but did not look at the stability of methylation after removing the stressor [34, 35]. One exception, Le Gac *et al.*, identified environment-related differentially methylated regions in poplar, but only examined stability across six months [36].

Detailed insights into the rate and spectrum of somatic mutations and epimutations are necessary to understand how somatic development of long-lived perennials contribute to population-level variation in an eco-evolutionary context. Here we generated a new high-quality reference genome from the oldest branch of a wild *Populus trichocarpa* tree with two dominant stems which have been evolving independently for approximately 330 years. By sampling multiple, age-estimated branches of this tree, we used a multi-omics approach to quantify age-related somatic changes at the genetic, epigenetic and transcriptional level. Our study provides the first quantitative insights into how nucleotide and functional variation arise during the lifetime of a long-lived perennial plant.

## RESULTS

### Experimental design for the discovery of somatic genetic and epigenetic variants

A stand of trees was identified near Mount Hood, Oregon and vegetative samples were collected from over 15 trees as part of an independent study. Of these trees, five were chosen for subsequent analysis and five branches of each tree were identified (Fig. S1). For each branch, the stem age was determined by coring the main stem at breast height and where the branch meets the stem and the branch age was determined by coring the base of the branch (Fig.1 and Fig. S2). Although 25 branches in total were initially sampled, six were excluded from analysis because they were epicormic and age estimates could not be determined. Two other branches had incomplete cores, but ages could be estimated based on radial diameter.

**Fig. 1.**
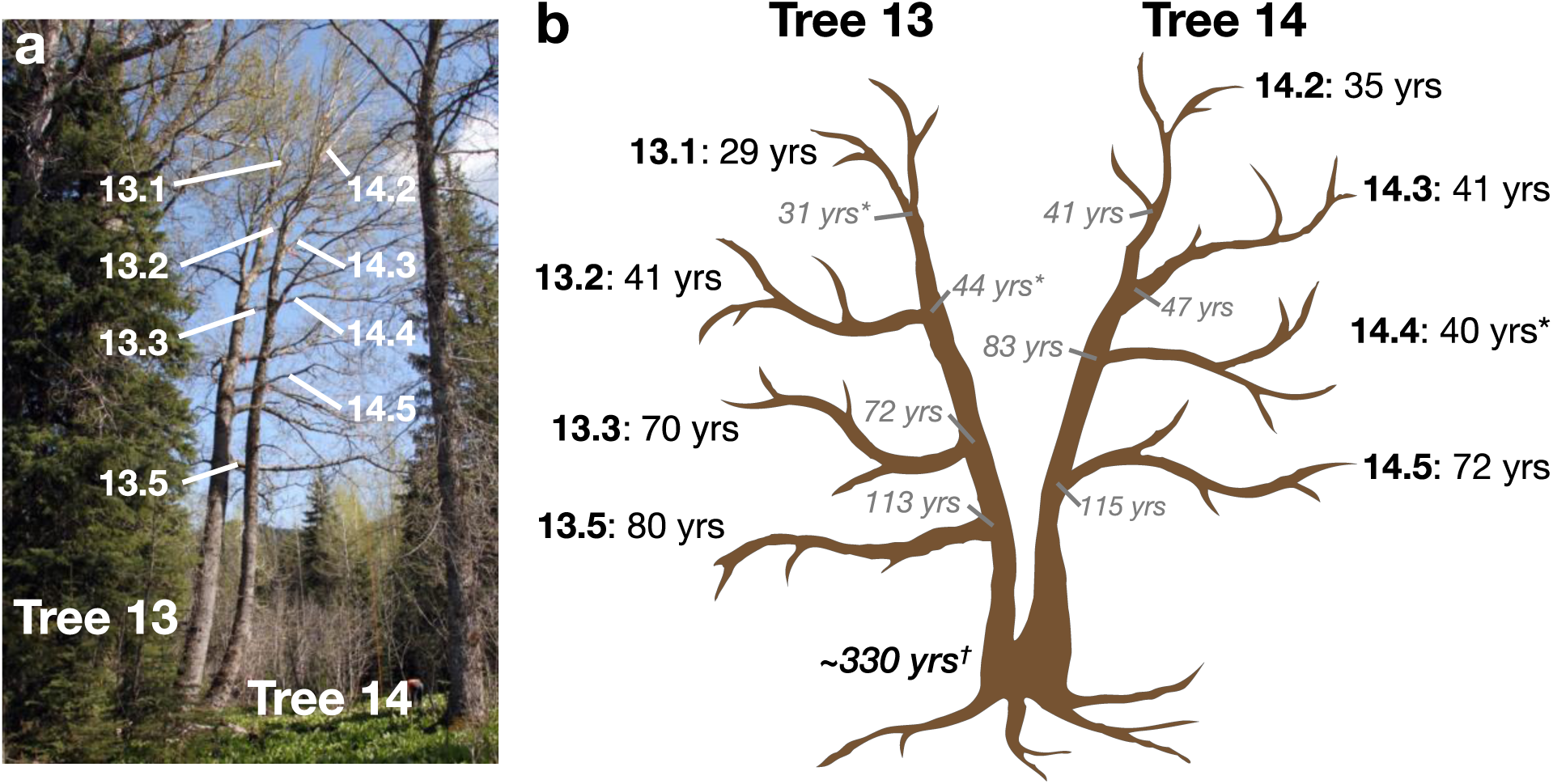
Photograph and schematic drawing of Tree 13 and Tree 14. This wild *P. trichocarpa*, located near Mt. Hood, Oregon, experienced a decapitation event ∼300 years ago. Tree 14 re-sprouted from the stump and ∼80-100 years later Tree 13 re-sprouted. (a) Leaf samples were collected from the labeled terminal branches. (b) Age was estimated for both the end of the branch (black font) and where it meets the main stem (gray italics). Ages with * indicate age was estimated using diameter; all other estimates were from core samples. Leaf samples of each branch was used to create genomic sequencing libraries, PacBio libraries, whole-genome bisulfite sequencing libraries, and mRNA-sequencing libraries.

From this, we were specifically interested in tree 13 and tree 14 (Fig. 1). Originally identified as two separate genotypes, they are actually two main stems of a single basal root system and trunk. Both tree 13 and tree 14 originated as stump sprouts off of an older tree that was knocked down over 300 years ago. Attempts to determine the total age were unsuccessful. However, statistical estimates based on molecular-clock arguments and a regression analysis of diameter to age suggest that the tree is approximately 330 years old (Shayary et al. 2019, co-submission).

Leaf samples were collected from eight age-estimated branches for multi-omics analysis for tree 13 and tree 14. The oldest branch of tree 14 (branch 14.5) was used for genome assembly of *Populus trichocarpa* var. *Stettler*. Genome resequencing was performed for all branches to explore intra- and inter-tree genetic variation. PacBio, MethylC-seq, and mRNA-seq libraries were constructed for the branches of tree 13 and tree 14 to explore structural, methylation, and transcriptional variation, respectively.

### Genome assembly and annotation of *Populus trichocarpa* var. *Stettler*

We sequenced the *P. trichocarpa* var. *Stettler* using a whole-genome shotgun sequencing strategy and standard sequencing protocols. Sequencing reads were collected using Illumina and PacBio. The current release is based on PacBio reads (average read length of 10,477 bp, average depth of 118.58x) assembled using the MECAT CANU v.1.4 assembler [37] and subsequently polished using QUIVER [38]. A set of 64,840 unique, non-repetitive, non-overlapping 1.0 kb sequences were identified in the version 4.0 *P. trichocarpa* var. *Nisqually* assembly and were used to assemble the chromosomes. The version 1 *Stettler* release contains 392.3 Mb of sequence with a contig N50 of 7.5 Mb and 99.8% of the assembled sequence captured in the chromosomes. Additionally, ∼232.2 Mb of alternative haplotypes were identified. Completeness of the final assembly was assessed using 35,172 annotated genes from the version 4.0 *P. trichocarpa* var. *Nisqually* release (jgi.doe.gov). A total of 34,327 (97.72%) aligned to the primary *Stettler* assembly.

The annotation was performed using ∼1.4 billion pairs of 2×150 stranded paired-end Illumina RNA-seq GeneAtlas *P. trichocarpa* var. *Nisqually* reads, ∼1.2 billion pairs of 2×100 paired-end Illumina RNA-seq *P. trichocarpa* var. *Nisqually* reads from Dr. Pankaj Jaiswal, and ∼430 million pairs of 2×75 stranded paired-end Illumina var. *Stettler* reads using PERTRAN (Shu, unpublished) on the *P. trichocarp*a var. *Stettler* genome. About ∼3 million PacBio Iso-Seq circular consensus sequences were corrected and collapsed by a genome-guided correction pipeline (Shu, unpublished) on the *P. trichocarpa* var. *Stettler* genome to obtain ∼0.5 million putative full-length transcripts. We annotated 34,700 protein-coding genes and 17,314 alternative splices for the final annotation. Because of the extensive resources included in the annotation, 32,330 genes had full-length transcript support.

### Identification and rate of somatic genetic variants

Leaf samples from the five trees were sequenced to an average depth of ∼87x (∼60-164x) using Illumina HiSeq. Roughly 88% of the high-quality reads map to the genome and about 98.6% of the genome is covered by at least one read, and genome coverage (∼8-500x) used for SNP calling was about 97%. The initial number of SNPs per tree (mutation on any branch) varied between 44,000 and 152,000, which is populated with many false positives due to coverage, sequencing and alignment errors, etc. Applying an additional filter requiring >20x coverage per position and requiring coverage in all branches reduced the total amount genome space queried to ∼40 Mb. Furthermore, since most of the genome (99.9%) is homozygous at every base pair, a somatic mutation will almost always result in a change from a homozygous to heterozygous site. Restricting the analysis to sites that change from homozygous to heterozygous, we identified 118 high-confidence SNPs in tree 13 and 143 high-confidence SNPs in tree 14 (Tables S1-2).

Over two-thirds of the SNPs in tree 13 and tree 14 were transition mutations, with C-G to T-A mutations accounting for over 54% of the SNPs (Fig. 2a). Of the transversion mutations C-G to G-C was the least common (3.8%) whereas C-G to A-T was most common (10%). Nearly half of the SNPs (46%) occurred in transposable elements and about 10% occur in promoter regions (Fig. 2b and Tables S1-S2). SNPs are significantly enriched in TEs and depleted in promoter regions genome-wide (Chi-square, df = 3, *P* < 0.001)

**Fig. 2.**
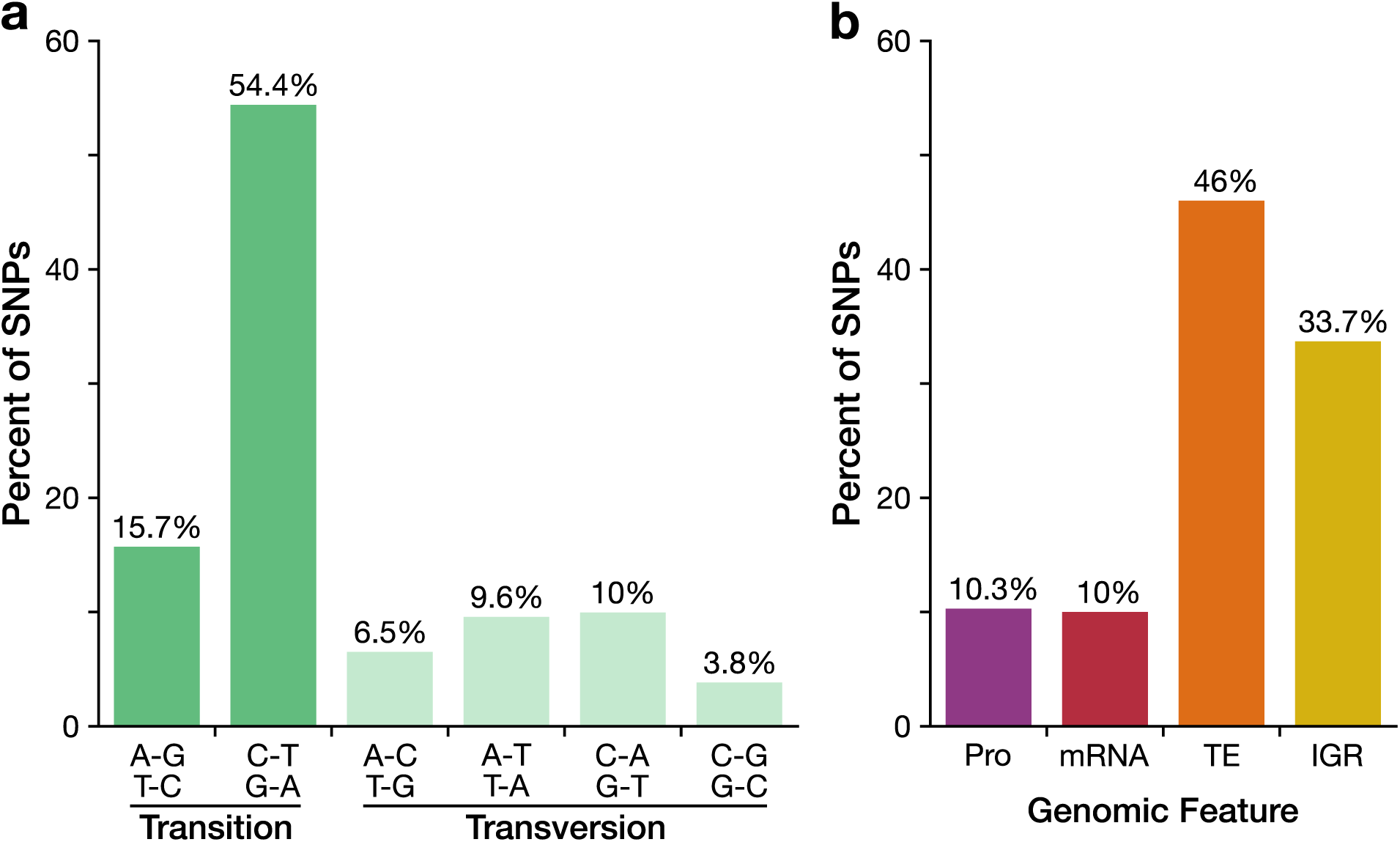
Most somatic mutations are transitions and occur in non-genic regions. (a) Distribution of reference to alternative allele observed in the high-confidence SNPs identified in Tree 13 and Tree 14. (b) Distribution of high-confidence SNPs separated by the genomic feature. Abbreviations: Pro, promoter; 2 kb upstream of TSS; TE, transposable elements and repeats; and IGR, intergenic regions.

To obtain an estimate of the rate of somatic point mutations from these SNP calls, we developed *mutSOMA* (https://github.com/jlab-code/mutSOMA), a phylogeny-based inference method that fully incorporates knowledge of the age-dated branching topology of the tree (see Methods and Supplementary Text). Using this approach, we find that the somatic point mutation rate in poplar is 1.33 × 10^−10^ (95% CI: 1.53 × 10^−11^ −4.18 × 10^−10^) per base per haploid genome per year (Table S3). Generation time can refer to two measurements—time from seed to production of first seeds and the organism’s lifespan. In annual plants, these values can be considered the same; however, this is not the case for perennials. Assuming 15 years from seed to production of first seeds [39], the poplar seed-to-seed generation mutation rate would be approximately 1.99 × 10^−9^. This is slightly lower than the per-generation (seed-to-seed) mutation rate observed in the annual *A. thaliana* (7 × 10^−9^) [16]. Next looking at the lifespan per-generation rate and assuming a maximum age of 200 years [40], the lifespan per-generation rate is 2.66 × 10^−8^. This estimate is slightly lower than the per-generation somatic mutation rate recently reported in oak (4.2 - 5.8 × 10^−8^) [14].

To analyze structural variants (SV) between haplotypes and somatic SV mutations, PacBio libraries were generated for the eight branches from tree 13 and tree 14 (Fig. 1). For each branch, four PacBio cells were sequenced generating an average output of 3.05 million reads and 28.3 Gb per branch (Table S4). After aligning the PacBio output to the *P. trichocarpa* var. *Stettler* genome, calling SVs larger than 20 bp, and filtering, we identified ∼10,466 deletions, ∼6,702 insertions, 645 duplications, and three inversions between the reference *Stettler* haplotype and the alternative haplotype (Table S5). Upon manual inspection of read mapping for a representative subset of SVs, 72.6% of SVs have strong support where multiple aligned reads support the SV type and size (Table S6). Deletions and duplications are significantly enriched in tandem repeat sequence and depleted in genic sequence (Kolmogorov-Smirnov two-sample test, *P* value < 2.2 × 10^−16^). Furthermore, deletions generally have less genic sequence and more tandem repeat sequence than do duplications (Fig. S3). Several of the detected SVs are large, with 11 deletions and five duplications greater than 50 kb (Table S5) with genic sequence content ranging from 0.0% to 23.7%. Comparisons of the branches from tree 13 and tree 14 did not identify instances of somatic SV mutation.

### Identification and rate of somatic epigenetic variants

To explore somatic epigenetic variation associated with changes in DNA methylation, we generated whole-genome bisulfite sequencing libraries from the branch tips of tree 13 and tree 14 (Fig. 1). The average genome coverage for the samples was ∼41.1x and sequence summary statistics are located in Table S7. Genome-wide methylation levels were similar across all samples with 36.61% mCG, 19.02% mCHG, and 2.07% mCHH% (Fig. S4) [41], indicating that global methylation levels are relatively stable across branches. Nonetheless, we observed significant age-dependent DNA methylation divergence between branches in CG and CHG contexts, both at the level of individual cytosines as well as at the level of regions, i.e. clusters of cytosines (Fig. 3a-b, Fig. S5, and Table S8). These age-dependent divergence patterns indicate that spontaneous methylation changes (i.e. epimutations) are cumulative across somatic development and thus point to a shared meristematic origin (Shahryary et al. 2019, co-submission).

**Fig. 3.**
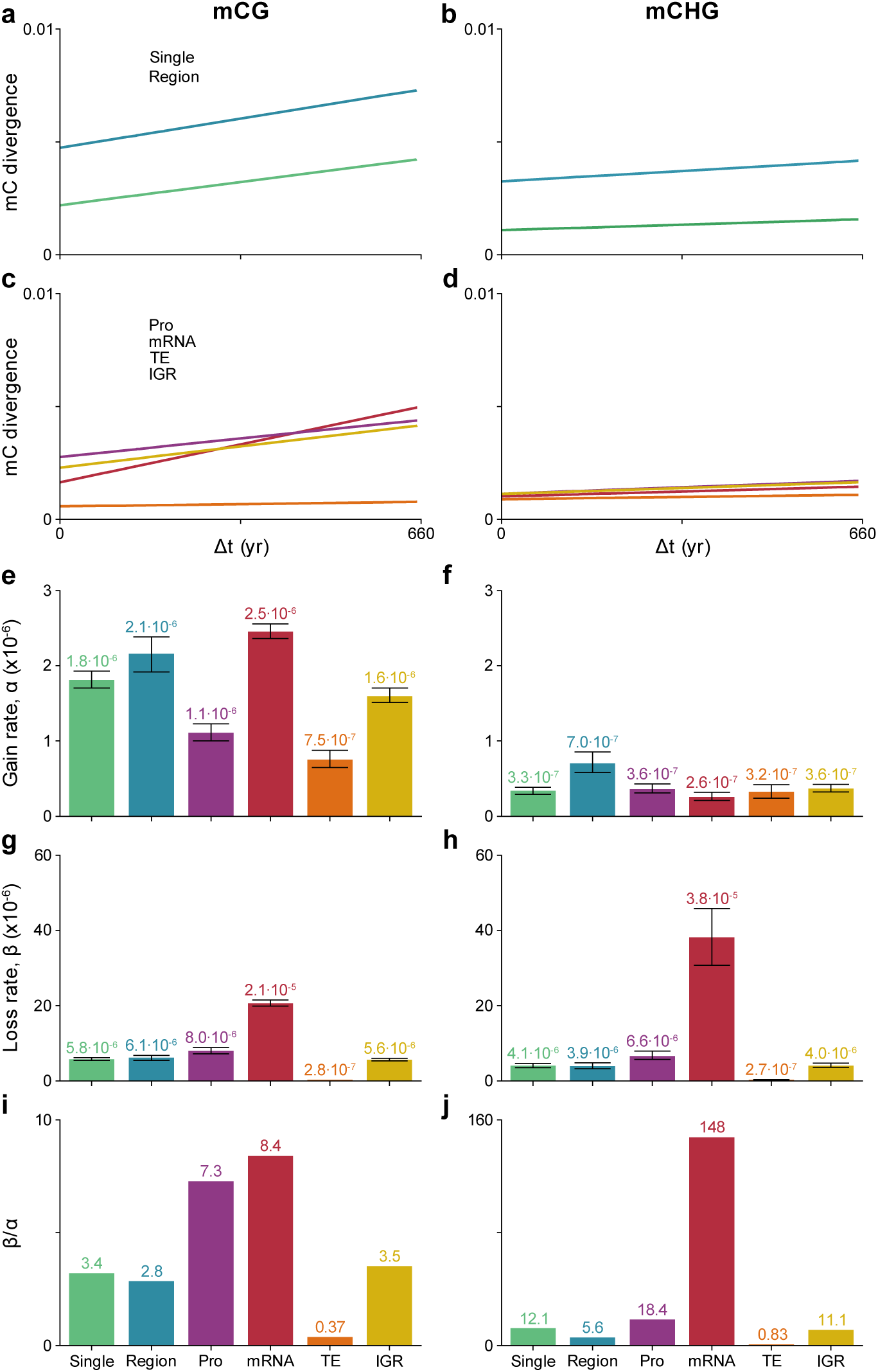
Somatic epimutation rates for single sites, regions, and by genomic feature. mCG (a) and mCHG (b) divergence by branch time divergence for single sites and regions; mCG (c) and mCHG (d) divergence by branch time divergence for genomic features Pro (promoter; 2 kb upstream of TSS), mRNA, TE (transposable elements), and IGR (intergenic regions); Estimated mCG (e) and mCHG (f) gain rates by feature; Estimated mCG (g) and mCHG (h) loss rates by feature; Ratio of mCG (i) and mCHG (j) loss to gain by feature. Error bars represent bootstrapped 95% confidence intervals of the estimates. Abbreviations: Pro, promoter; 1.5 kb upstream of TSS; TE, transposable elements and repeats; and IGR, intergenic regions.

To obtain an estimate of somatic epimutation rates, we applied *AlphaBeta* (Shahryary et al. 2019, co-submission). The method builds on our previous approach for estimating ‘germline’-epimutation in mutation accumulation (MA) lines [32], except here we treat the tree branching topology as an intra-organismal phylogeny and model mitotic instead of meiotic inheritance. Focusing first on cytosine-level epimutations, we estimated that at the genome-wide scale spontaneous methylation gains in contexts CG and CHG occur at a rate of 1.8 × 10^−6^ and 3.3 × 10^−7^ per site per haploid genome per year, respectively; whereas spontaneous methylation losses in these two sequence contexts occur at a rate of 5.8 × 10^−6^ and 4.1 × 10^−6^ per site per haploid genome per year. Based on these estimates, we extrapolate that the *seed-to-seed* per-generation epimutation rate in poplar is about 10-5 and the *lifespan* per-generation rate is 10-4. Remarkably, these estimates are very similar to those reported in *A. thaliana* MA lines [32]. The observation that two species with such different life history traits and genome architecture display very similar per-generation mutation and epimutation rates suggests that the rates themselves are subject to strong evolutionary constraints.

In addition to global epimutation rates, we also estimated rates for different genomic features (mRNA, promoters, intergenic, TEs). This analysis revealed highly significant rate differences in the CG and CHG context between genomic features, with mRNAs showing the highest and TEs the lowest combined rates (Fig. 3c-j). Interestingly, the ordering of the magnitude of the mRNA, promoter, and intergenic rates is similar to that previously observed in *A. thaliana* MA lines [32]. The differences in rates at local genomic features likely reflect the distinct DNA methylation pathways that function on these sequences (RNA-directed DNA methylation, CHROMOMETHYLASE3, CHROMOMETHYLASE2, DNA METHYLTRANSFERASE1, etc.). For example, the high rate of epimutation losses in mRNA relative to other features (Fig. 3g-h) could reflect the activity of CMT3-mediated gene body DNA methylation [42, 43]. The observation that the epimutation rates of these features is consistent between *A. thaliana* MA lines (>60 generations) and this long-lived perennial (within a single generation) seems to imply that epimutations are not a result of biased reinforcement of DNA methylation during sexual reproduction or environment/genetic variation, but instead a feature of DNA methylation maintenance through mitotic cell divisions.

### Assessment of spontaneous differentially methylated regions

Differentially methylated regions are functionally more relevant than individual cytosine-level changes, as in certain cases they are linked to differential gene expression and phenotypic variation [26–28, 44, 45]. To explore the extent of differentially methylated regions (DMRs) that spontaneously arise in these trees we searched for all pairwise DMRs between all branches. In total, we identified 10,909 DMRs that possessed changes in all sequence contexts (CG, CHG and CHH - C-DMRs). Together they constitute approximately 1.69 Mb of the total 167.4 Mb (∼1%) of methylated sequences in the *Stettler* genome and they reveal age-dependent accumulation (Fig. 4a). Most DMRs occur in intergenic regions (56.7%), but a significant enrichment of DMRs were detected within two kilobases from the transcriptional start site of genes compared to methylated regions as a whole (Fig. 4b) (Fisher’s exact test, one-sided, *P* value < 0.001).

**Fig. 4.**
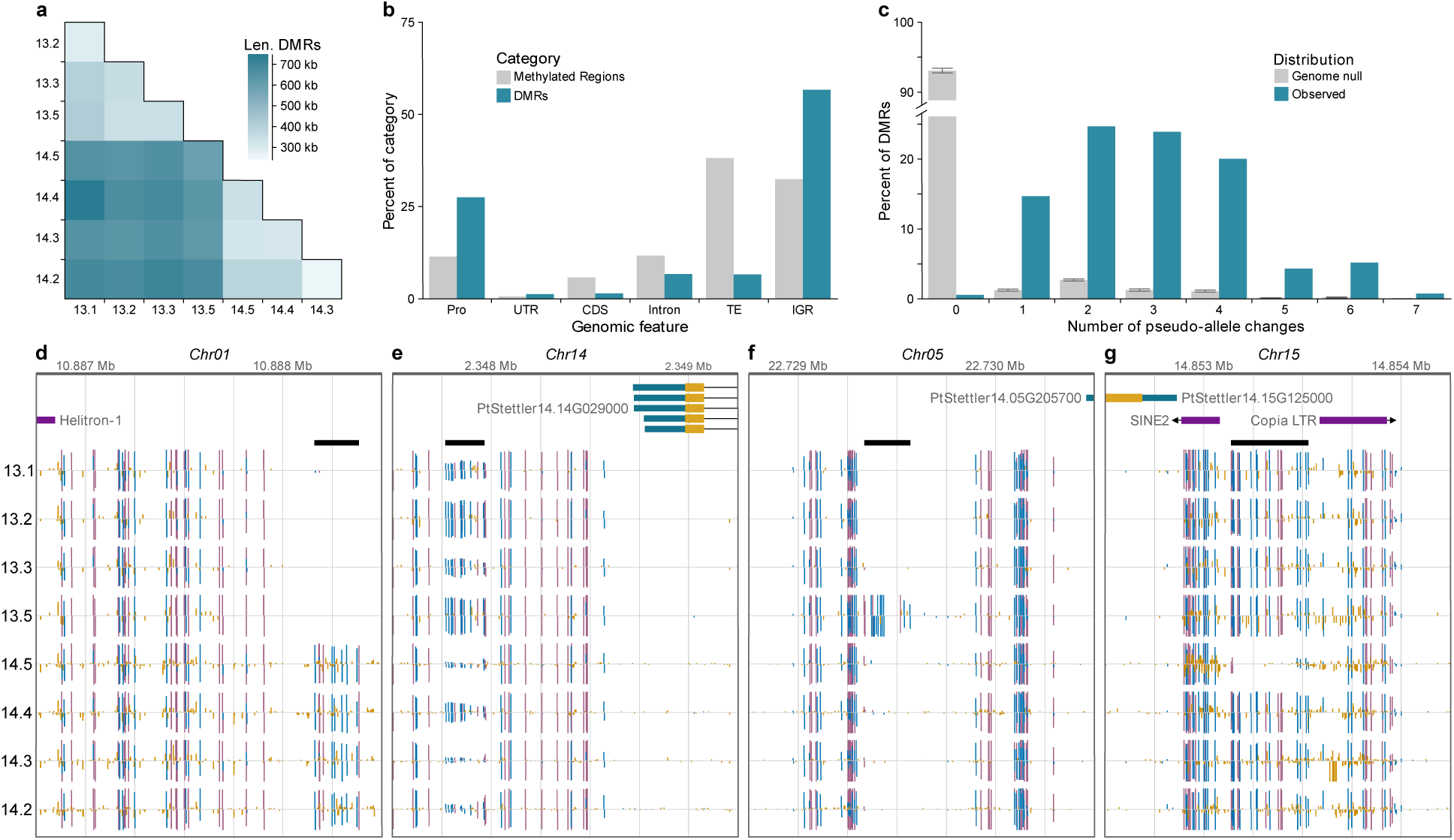
Identification and quantification of somatic stability of differentially methylated regions. (a) Divergence of differentially methylated regions corresponds to divergence in age. The darker color indicates combined length of the pairwise DMRs; (b) The genome-wide distribution of identified DMRs in genomic features. Abbreviations: TE, transposable elements and repeats; IGR, intergenic region; Pro, promoter region (2 kb upstream of transcription start site); UTR, untranslated regions; CDS, coding sequence. Methylated regions were identified in as regions methylated in at least one sample. (c) There are significantly more pseudo-allele changes between the branches at DMRs (blue) compared to the genome-wide null (Wilcox rank sum, one-sided, *P* value < 2 × 10^−16^). Gray bars are the genome-wide null as mean +/- std. dev. across 10 simulations. (d) Browser screenshot of a tree specific DMR where all branches in tree 13 are homozygous unmethylated and all branches of tree 14 are homozygous methylated. (e) Browser screenshot of a highly variable DMR where the pseudo allele state changes between each branch. (f) Browser screenshot of a single gain DMR where all branches except 13.5 are homozygous unmethylated and 13.5 gains methylation. (g) Browser screenshot of a single loss DMR where all branches except 14.5 are homozygous methylated and 14.5 has lost methylation. For d-g, gene models and transposable elements are shown at the top and methylome tracks are below. Vertical bars indicate methylation at the position, where height corresponds to level and color is context, red for CG, blue for CHG, and yellow for CHH. DMR is indicated by thick black horizontal line.

Given the heterozygous nature of wild *P. trichocarpa*, we explored allelic methylation changes. After filtering for sufficient coverage and methylation change, we assigned the pseudo-allele state of each branch at 4,488 DMRs. Possible states were homozygous unmethylated, heterozygous, and homozygous methylated. In each sample, 43.0% of DMRs, on average, were categorized as homozygous methylated (Fig. S6). Interestingly, the youngest branches, 13.1 and 14.1 have about 10% more homozygous methylated pseudo-alleles than the other branches (51.1% vs 41.7%). Next, we looked at the number of changes of pseudo-allele states. This is expected as DMRs were identified as having different methylation levels in the samples. On average, there are 3.02 state changes for each DMR with 94.4% of DMRs having one to five state changes (Fig. 4c). These data suggest that many of these regions are metastable, a common feature of epimutations in plants.

An example of a region with one state change are the tree specific DMRs (Fig. 4d). In these regions, all branches of one tree are homozygous unmethylated and all branches of the other tree are homozygous methylated. This suggesting the methylation state change occurred shortly after the trees separated and remained stable throughout subsequent mitotic divisions. In contrast, we also identified highly variable regions with seven state changes, a change between each branch (Fig. 4e). Of the regions with two state changes, 150 have branch-specific state changes. For example, in Fig. 4f branches 13.1 to 13.3 are homozygous unmethylated, then it changes to homozygous methylated for branch 13.5, and changes again to homozygous unmethylated for branches 14.5 – 14.2. Similarly, in Fig. 4g, all branches except 14.5 are homozygous methylated and 14.5 has spontaneously lost methylation.

We also used the identified C-DMRs (differential methylation in all cytosine sequence contexts) to obtain region-level epimutation rates. To do this, we established control regions (‘non-DMR’) with the same size distribution as observed for C-DMRs and used the methylation levels of all cytosines in each (non-)DMR to calculate methylation levels per region. Interestingly, this analysis shows that region-level epimutation rates are comparable to epimutation rates of single cytosines. Even though there are far fewer DMRs in comparison to epimutations at single cytosines, the similar rates are not too unexpected if one considers that the total ‘epimutable space’ for regions in the genome is much smaller than that for individual cytosines. In summary, these results might suggest that the mechanisms which underlie spontaneous differential methylation are the same for differential methylation in larger regions and at individual sites.

### Functional consequences of differential methylation on gene expression

To assess if age-related cytosine methylation changes have functional consequences, we performed mRNA-seq with three biological replicates for each branch of trees 13 and 14. On average, each library had over ∼55 million reads and 96.8% mapping to the *P. trichocarpa* var. *Stettler* genome (Table S9). We used DESeq2 to identify differentially expressed genes (DEGs) pairwise between branches [46] and identified a total of 2,937 genes. The *P. trichocarpa* var. *Stettler* genome has 34,700 annotated genes, so this differential expression gene set is 8.46% of all genes and 10.5% of expressed genes.

Since the somatic accumulation of spontaneous methylation changes could affect gene expression, we asked if transcriptional divergence also increases as a function of tree age. We found that in contrast to somatic mutations and epimutations, the divergence between leaf transcriptomes is much more heterogeneous and displays only a weak and non-significant accumulation trend (Fig. 5a). This observation suggests that the accumulation of genetic and epigenetic changes are largely uncoupled from age-dependent transcriptional changes in poplar, at least at the global scale.

**Fig. 5.**
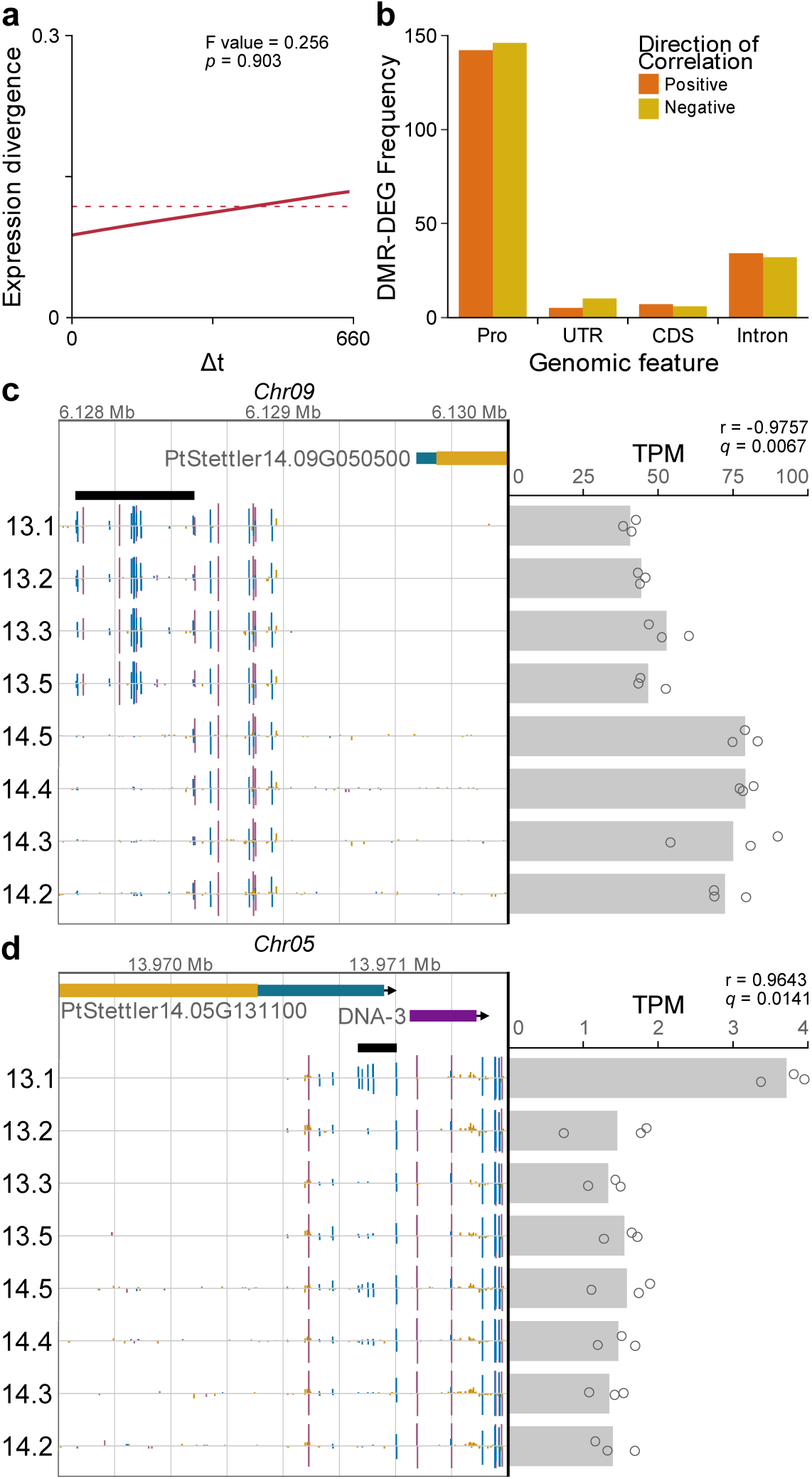
Gene expression is largely independent from divergence age and nearby cytosine methylation except in a few examples. a) Gene expression divergence is not significantly associated with divergence age. b) Distribution of positive and negative correlations for differentially expressed genes and overlapping/nearby DMRs. Positive correlation occurs when the higher methylation level is associated with higher gene expression among the samples. (c) A significantly negatively correlated, tree-specific DMR and DEG where the DMR occurs in the promoter region of the gene (Pearson’s correlation test, two-sided, N = 8, adjusted *P* value = 0.0067). The higher methylation levels in the DMR for tree 13 branches are associated with lower gene expression. (d) A significantly positively correlated, single gain DMR and DEG where the DMR occurs in the 5’ untranslated region of the gene (Pearson’s correlation test, two-sided, N = 8, adjusted *P* = 0.0141). The higher methylation level in the DMR for branch 13.1 is associated with greater gene expression. For c and d, gene expression, as transcripts per million (TPM), is represented as points for the individual replicates and as bar for mean among replicates. In the genome browser view, gene models and transposable elements are shown at the top and methylome tracks are below. Vertical bars indicate methylation at the position, where height corresponds to level and color is context, red for CG, blue for CHG, and yellow for CHH. DMR is indicated by thick black horizontal line.

However, this global analysis does not rule out that DNA methylation changes at specific individual loci can have transcriptional consequences. To explore this in more detail, we analyzed DMRs proximal to DEGs, and correlated the methylation level of the DMR with the expression level of the gene. The correlation is positive when a higher methylation level in the DMR is associated with higher expression of the gene.

Regardless of where the DMR was located relative to the gene, we observed positive DMR-DEG correlations and negative DMR-DEG correlations. There was no bias for direction of correlation and genomic feature type (Fig. 5b).

We further focused on four specific examples where DEG-DMR correlations were statistically significant (Fig. S7). Of these four, three of the DMRs occurred within two kilobases upstream of the transcription start site, and they have strong negative correlations (Fig. 5c). The DMR located in the untranslated region of a gene encoding a mitochondrial oxoglutarate/malate carrier protein was positively correlated with gene expression (Fig. 5d), although it remains unclear if this relationship is causal.

Taken together, our transcriptome analysis indicates that gene expression changes in this poplar tree are largely independent of methylation at both the global and local scale except for a few rare examples. This observation is at least partly consistent with our model-based analyses, which suggest that somatic epimutations in this tree accumulate neutrally (Shahryary et al. 2019, co-submission).

## DISCUSSION

Using a multi-omics approach, we were able to calculate the rates of somatic mutations and epimutations in the long-lived perennial tree *P. trichocarpa*. Consistent with the per-unit-time hypothesis, we find that the per-year genetic and epigenetic mutation rates in poplar are lower than in A. thaliana, which is remarkable considering that the former experienced hundreds of years of variable environmental conditions. This observation supports the view that long-lived perennials may limit the number of meristematic cell divisions during their lifetime and that they have evolved mechanisms to protect these cell types from the persistent influence of environmental mutagens, such as UV-radiation. Interestingly, in contrast to the observed differences in *per-year* mutation and epimutation rates, our analysis reveals strong similarities in the *per-generation* rates between these two species. This close similarity further suggests that the per-generation rates of spontaneous genetic and epigenetic changes are under strong evolution constraint, although it remains unclear from our experimental design how many of these (epi)mutations will be successfully transferred to the next generation.

The results presented here are most certainly an underestimate of the actual rate. This may be a result of the sampling biased used in this study, as we were only able to sample surviving branches and identify mutations that occurred early enough that they are present in the majority of the cells sampled in the tissues profiled. Perhaps variable environmental conditions lower the epimutation rate by keeping the cells in sync, thus few differences can be observed. Alternatively, meristematic cells that give rise to the sampled tissues have highly reinforced and well-maintained DNA methylomes similar to observations in embryonic tissue [47–51]. Either scenario would imply that most of the identified epimutations are spontaneous in nature. Although the rate is different, the ordering in feature-specific epimutation rates is the same between poplar and *A. thaliana*, suggesting that this is a general pattern in plant genomes, which likely is derived from maintenance of DNA methylation through mitotic cell divisions.

## CONCLUSION

Taken together, our study provides unprecedented insights into the origin of nucleotide, epigenetic, and functional variation in the long-lived perennial plant.

## METHODS

### Sample collection and age estimation

The trees used in this study were located at Hood River Ranger District [Horse Thief Meadows area], Mt. Hood National Forest, 0.6 mi south of Nottingham Campground off OR-35 at unmarked parking area, 500’ west of East Fork Trail #650 across river, ca. 45.355313, −121.574284 (Fig. S1).

Cores were originally collected from the main stem and five branches from each of five trees in April 2015 at breast height (∼1.5 m) for standing tree age using a stainless-steel increment borer (5 mm in diameter and up to 28 cm in length). Cores were mounted on grooved wood trim, dried at room temperature, sanded and stained with 1% phloroglucinol following the manufacturer’s instructions (https://www.forestry-suppliers.com/Documents/1568_msds.pdf). Annual growth rings were counted to estimate age. For cores for which accurate estimates could not be made from the 2015 collection, additional collections were made in spring 2016. However, due to difficulty in collecting by climbing, many of the cores did not reach the center of the stem or branches (pith) and/or the samples suffered from heart rot. Combined with the difficulty in demarcating rings in porous woods such as poplar *Populus* [52, 53], accurate measures of tree age or branch age were challenging (Fig. S2).

Simultaneously with stem coring, leaf samples were collected from the tips of each of the branches from the selected five trees. Branches 9.1, 9.5, 13.4, 14.1, 15.1, and 15.5 were too damaged to determine reasonable age estimates and were removed from analysis. Branch 14.4 and the stems of 13.1 and 13.2 were estimated by simply regressing the diameter of all branches and stems that could be aged by coring.

### Nuclei prep for DNA extraction

Poplar leaves, that had been kept frozen at −80 °C, were gently ground with liquid nitrogen and incubated with NIB buffer (10 mM Tris-HCL, PH8.0, 10 mM EDTA PH8.0, 100 mM KCL, 0.5 M sucrose, 4 mM spermidine, 1 mM spermine) on ice for 15 min. After filtration through miracloth, Triton x-100 (Sigma) was added to tubes at a 1:20 ratio, placed on ice for 15 min, and centrifuged to collect nuclei. Nuclei were washed with NIB buffer (containing Triton x-100) and re-suspended in a small amount of NIB buffer (containing Triton x-100) then the volume of each tube was brought to 40 ml and centrifuged again. After careful removal of all liquid, 10 ml of Qiagen G2 buffer was added followed by gentle re-suspension of nuclei; then 30 ml G2 buffer with RNase A (to final concentration of 50 mg/ml) was added. Tubes were incubated at 37 °C for 30 min. Proteinase K (Invitrogen), 30 mg, was added and tubes were incubated at 50 °C for 2 h followed by centrifugation for 15 min at 8000 rpm, at 4 °C, and the liquid gently poured to a new tube. After gentle extraction with Chloroform / isoamyl alcohol (24:1), then centrifugation and transfer of the top phase to a fresh tube, HMW DNA was precipitated by addition of 2/3 volume of iso-propanol and re-centrifugation to collect the DNA. After DNA was washed with 70% ethanol, it was air dried for 20 min and dissolved thoroughly in 1x TE.

### Whole-genome sequencing

We sequenced *Populus trichocarpa* var. *Stettler* using a whole-genome shotgun sequencing strategy and standard sequencing protocols. Sequencing reads were collected using Illumina and PacBio. Both the Illumina and PacBio reads were sequenced at the Department of Energy (DOE) Joint Genome Institute (JGI) in Walnut Creek, California and the HudsonAlpha Institute in Huntsville, Alabama. Illumina reads were sequenced using the Illumina HISeq platform, while the PacBio reads were sequenced using the RS platform. One 400-bp insert 2×150 Illumina fragment library was obtained for a total of ∼349x coverage (Table S10). Prior to assembly, all Illumina reads were screened for mitochondria, chloroplast, and phix contamination. Reads composed of >95% simple sequence were removed. Illumina reads less than 75 bp after trimming for adapter and quality (q < 20) were removed. The final Illumina read set consists of 906,280,916 reads for a total of ∼349x of high-quality Illumina bases. For the PacBio sequencing, a total of 69 chips (P6C4 chemistry) were sequenced with a total yield of 59.29 Gb (118.58x) with 56.2 Gb > 5 kb (Table S11), and post error correction a total of 37.3 Gb (53.4x) was used in the assembly.

### Genome assembly and construction of pseudomolecule chromosomes

The current release is version 1.0 release began by assembling the 37.3 Gb corrected PacBio reads (53.4x sequence coverage) using the MECAT CANU v.1.4 assembler [37] and subsequently polished using QUIVER v.2.3.3 [38]. This produced 3,693 scaffolds (3,693 contigs), with a scaffold N50 of 1.9 Mb, 955 scaffolds larger than 100 kb, and a total genome size of 693.8 Mb (Table S12). Alternative haplotypes were identified in the initial assembly using an in-house Python pipeline, resulting in 2,972 contigs (232.3 Mb) being labeled as alternative haplotypes, leaving 745 contigs (461.5 Mb) in the single haplotype assembly. A set of 64,840 unique, non-repetitive, non-overlapping 1.0 kb syntenic sequences from version 4.0 *P. trichocarpa* var. *Nisqually* assembly and aligned to the MECAT CANU v.1.4 assembly and used to identify misjoins in the *P. trichocarpa* var. *Stettler* assembly. A total of 22 misjoins were identified and broken. Scaffolds were then oriented, ordered, and joined together into 19 chromosomes. A total of 117 joins were made during this process, and the chromosome joins were padded with 10,000 Ns. Small adjacent alternative haplotypes were identified on the joined contig set. Althap regions were collapsed using the longest common substring between the two haplotypes. A total of 14 adjacent alternative haplotypes were collapsed.

The resulting assembly was then screened for contamination. Homozygous single nucleotide polymorphisms (SNPs) and insertion/deletions (InDels) were corrected in the release sequence using ∼100x of Illumina reads (2×150, 400-bp insert) by aligning the reads using bwa-0.7.17 mem [54] and identifying homozygous SNPs and InDels with the GATK v3.6’s UnifiedGenotyper tool [55]. A total of 206 homozygous SNPs and 11,220 homozygous InDels were corrected in the release. Heterozygous SNP/indel phasing errors were corrected in the consensus using the 118.58x raw PacBio data. A total of 66,124 (1.98%) of the heterozygous SNP/InDels were corrected. The final version 1.0 improved release contains 391.2 Mb of sequence, consisting of 25 scaffolds (128 contigs) with a contig N50 of 7.5 Mb and a total of 99.8% of assembled bases in chromosomes. Plots of the *Nisqually* marker placements for the 19 chromosomes are shown in Fig. S8.

### Genome annotation

Transcript assemblies were made from ∼1.4 billion pairs of 2×150 stranded paired-end Illumina RNA-seq GeneAtlas *P. trichocarpa* Nisqually reads, ∼1.2 billion pairs of 2×100 paired-end Illumina RNA-seq *P. trichocarpa* Nisqually reads from Dr. Pankaj Jaiswal, and ∼430M pairs of 2×75 stranded paired-end Illumina var. *Stettler* reads using PERTRAN (Shu, unpublished) on *P. trichocarpa* var. *Stettler* genome. About ∼3M PacBio Iso-Seq circular consensus sequences were corrected and collapsed by genome guided correction pipeline (Shu, unpublished) on *P. trichocarpa* var. *Stettler* genome to obtain ∼0.5 million putative full-length transcripts. 293,637 transcript assemblies were constructed using PASA [56] from RNA-seq transcript assemblies above. Loci were determined by transcript assembly alignments and/or EXONERATE alignments of proteins from *A. thaliana*, soybean, peach, Kitaake rice, *Setaria viridis*, tomato, cassava, grape and Swiss-Prot proteomes to repeat-soft-masked *P. trichocarpa* var. *Stettler* genome using RepeatMasker [57] with up to 2-kb extension on both ends unless extending into another locus on the same strand. Gene models were predicted by homology-based predictors, FGENESH+[58], FGENESH_EST (similar to FGENESH+, EST as splice site and intron input instead of protein/translated ORF), and EXONERATE [59], PASA assembly ORFs (in-house homology constrained ORF finder) and from AUGUSTUS via BRAKER1 [60]. The best scored predictions for each locus are selected using multiple positive factors including EST and protein support, and one negative factor: overlap with repeats. The selected gene predictions were improved by PASA. Improvement includes adding UTRs, splicing correction, and adding alternative transcripts. PASA-improved gene model proteins were subject to protein homology analysis to above mentioned proteomes to obtain Cscore and protein coverage. Cscore is a protein BLASTP score ratio to MBH (mutual best hit) BLASTP score and protein coverage is highest percentage of protein aligned to the best of homologs. PASA-improved transcripts were selected based on Cscore, protein coverage, EST coverage, and its CDS overlapping with repeats. The transcripts were selected if its Cscore is larger than or equal to 0.5 and protein coverage larger than or equal to 0.5, or it has EST coverage, but its CDS overlapping with repeats is less than 20%. For gene models whose CDS overlaps with repeats for more that 20%, its Cscore must be at least 0.9 and homology coverage at least 70% to be selected. The selected gene models were subject to Pfam analysis and gene models whose protein is more than 30% in Pfam TE domains were removed and weak gene models. Incomplete gene models, low homology supported without fully transcriptome supported gene models and short single exon (< 300-bp CDS) without protein domain nor good expression gene models were manually filtered out.

### SNP calling methods

Illumina HiSeq2500 paired-end (2×150) reads were mapped to the reference genome using bwa-mem [54]. Picard toolkit was used to sort and index the bam files. GATK [55] was used further to align regions around InDels. Samtools v1.9 [61] was used to create a multi-sample mileup for each tree independently. Preliminary SNPs were called using Varscan v2.4.0 [62] with a minimum coverage of 21.

At these SNPs, for each branch, we calculated the conditional probability of each potential genotype (RR, RA, AA) given the read counts of each allele, following SeqEM [63], using an estimated sequencing error rate of 0.01. We identified high-confidence genotype calls as those with a conditional probability 10,000x greater than the probabilities of the other possible genotypes. We identified potential somatic SNPs as those with both a high-confidence homozygous and high-confidence heterozygous genotype across the branches.

We notice that the default SNP calling parameters tend to overcall homozygous-reference allele genotypes and that differences in sequencing depth can bias the relative number of heterozygous SNPs detected. To overcome these issues, we re-called genotypes using conditional probabilities using down sampled allele counts. To do this, we first randomly selected a set number of sequencing reads for each library at each potential somatic SNP so that all libraries have the same sequencing depth at all SNPs. Using the down sampled reads, we calculate the relative conditional probability of each genotypes by dividing the conditional probabilities by the sum of the conditional probabilities of all three potential genotypes. These relative probabilities are then multiplied by the dosage assigned to their respective genotype (0 for RR, 1 for RA, 2 for AA), and the dosage genotype is the sum of these values across all 3 possible genotypes. Discrete genotypes were assigned using the following dosage values: RR = dosage < 0.1; RA = 0.9 < dosage < 1.1; AA = dosage > 1.9. Dosages outside those ranges are assigned a NA discrete genotype. SNPs with an NA discrete genotype or depth below the down sampling level in any branch of a tree were removed from further analysis. We performed three replicates of this procedure for depths of 20, 25, 30, 35, 40, and 45 reads.

PacBio libraries for each branch were sequenced using the PacBio Sequel platform, fastq files aligned to the *P. trichocarpa* var. Stettler14 reference genome using ngmlr [64], and multi-sample mileup files generated using in Samtools v1.9 [61] to quantify the allele counts at the potential somatic SNPs. We used a minimum per-sample sequence depth of 20 reads and used an alternate-allele threshold of 0.1 to call a heterozygote genotype in the PacBio data.

To identify high-confidence candidate somatic SNPs, we identified potential somatic SNPs with the same genotypes across branches using both the Illumina-based PacBio-based genotypes, only including SNPs with full data in all branches for both types of genotypes. Of these, we only retained SNPs that are homozygous in a single branch or have a single homozygous-to-heterozygous transition (and no reversion) going from the lowest to highest branches.

### Estimating somatic nucleotide mutation rate

Building on the analytical framework developed in van der Graaf et al. (2015) and Shahryary et al. 2019 (co-submission), we developed *mutSOMA* (https://github.com/jlab-code/mutSOMA), a statistical method for estimating genetic mutation rates in long-lived perennials such as trees. The method treats the tree branching structure as a pedigree of somatic lineages and uses the fact that these cell lineages carry information about the mutational history of each branch. A detailed mathematical description of the method is provided in Supplementary Text. But briefly, starting from the .vcf* files from *S* samples representing different branches of the tree, we let *G_ik_* be the observed genotype at the *k-*th single nucleotide (*k* = 1, …, *N*) in the *i*-th sample, where *N* is the effective genome size (i.e. the total number of bases with sufficient coverage). With four possible nucleotides (A, C, T, G, G_ik_ can have 16 possible genotypes in a diploid genome, 4 homozygous (A|A, T|T, C|C, G|G) and 12 heterozygous (A|G, A|T, …, G|C). Using this coding, we calculate the genetic divergence, *D*, between any two samples *i* and *j* as follows:

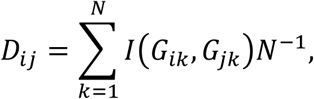

where *I*(*G_ik_*, *G_jk_*) is an indicator function, such that, *I*(*G_ik_*, *G_jk_*) = 1 if the two samples share no alleles at locus *k*, 0.5 if they share one, and 0 if they share both alleles. We suppose that *D_ij_* is related to the developmental divergence time of samples *i* and *j* through a somatic mutation model *M_Θ_* . The divergence times can be calculated from the coring data (Table S13). We model the genetic divergence using

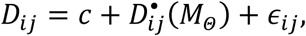

where *ϵ_ij_* ∼ *N*(0, *σ*^2^) is the normally distributed residual, *c* is the intercept, and 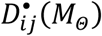 is the expected divergence as a function of mutation model *M* with parameter vector ϴ. Parameter vector ϴ contains the unknown mutation rate *δ* and the unknown proportion *γ* heterozygote loci of the most recent common ‘founder’ cells of samples *i* and *j*. The theoretical derivation of 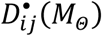 and details regarding model estimation can be found in Supplementary Text. The estimation of the residual variance in the model allows for the fact that part of the observed genetic divergence between any two samples is driven both by genotyping errors as well as by somatic genetic drift as meristematic cells pass through bottlenecks in the generation of the lateral branches.

### Structural variant analysis methods

For structural variant (SV) analysis, PacBio libraries were generated for four branches from the tree 13 and four branches from tree 14 with four sequencing cells sequenced per branch using the PacBio Sequel platform. PacBio fastq files were aligned to the *P. trichocarpa* var. *Stettler* reference genome using ngmlr v.0.2.6 [64] using a value of 0.01 for the “-R” flag. SVs were discovered and called using pbsv (pbsv v2.2.0, https://github.com/PacificBiosciences/pbsv). SV signatures were identified for each sample using ‘pbsv discover’ using the ‘--tandem-repeats’ flag and a tandem repeat BED file generated using trf v4.09 [65] for the *P. trichocarpa* var. *Stettler* genome. SVs were called jointly for all 8 branches using ‘pbsv call’. The output from joint SV calling changes slightly depending on the order of the samples used for the input in ‘pbsv call’, so four sets of SVs were generated using four different sample orders as input. We used a custom R script [66] to filter the SV output from pbsv. We remove low-complexity insertions or deletions with sequence containing > 80% of a mononucleotide 8-mer, 50% of a single type of binucleotide 8-mer, or 60% of two types of binucleotide 8-mers. We required a minimum distance of 1 kb between SVs. We removed SVs with sequencing coverage of more than three standard deviations above the mean coverage across a sample. After calling genotypes, any SVs with missing genotype data were removed.

Genotypes were called based on the output from pbsv using a custom R script. We required a minimum coverage of 10 reads in all sample and for one sample to have at least 20 reads. We required a minimum penetrance (read ratio) of 0.25 and at least 2 reads containing the minor allele for a heterozygous genotype. We allowed a maximum penetrance of 0.05 for homozygous genotypes. For each genotype, we assigned a quality score based on the binomial distribution-related relative probability of the 3 genotype classes (RR, AR, AA) based on A:R read ratio, using an estimated sequencing error of 0.032, and an estimated minimum allele penetrance of 0.35. For a genotype with a score below 0.9 but with the same genotype at the SV as another sample with a score above 0.98, the score was adjusted by multiplying by 1.67. Any genotypes with adjusted scores below 0.9 were converted to NA. For deletions, duplications, and insertions, 10 representatives in different size classes were randomly selected and the mapping patterns of reads were visually inspected using IGV v2.5.3 [67] to assign scores indicating how well the visual mapping patterns support the SV designation. Scores were defined by the following: “strong”, multiple reads align to the same locations in the reference genome that support the SV type and size; “moderate”, multiple reads align to the same reference location for one side of the SV but align to different or multiple locations in the region for the other side of the SV; and “weak”, reads align to reference locations that indicate a different SV type or much different SV size.

The percent of genic sequence and tandem repeat sequence in deletions and duplications were calculated using the *P. trichocarpa* var. *Stettler* annotation and tandem repeat BED from above, respectively. Genome-wide expectations were derived by separating the genome into 10-kb windows and calculating the percent genic and tandem repeat sequence in each window. The distribution of genic and tandem repeat sequences in deletions and duplications were compared to genome-wide expectations using the Kolmogorov-Smirnov two-sample test (one-sided, N_null_ = 39,151, N_del_ = 10,433, N_dup_ = 630).

SVs showing variation between branches and identified in all 4 replicates are potential instances of somatic SV mutations or loss-of-heterozygosity gene conversions, and the mapping positions of sequencing reads were visually inspected with IGV [67] to confirm the variation at these SVs.

### MethylC-seq sequencing and analysis

A single MethylC-seq library was created for each branch from leaf tissue. Libraries were prepared according to the protocol described in Urich *et al.* [68]. Libraries were sequenced to 150-bp per read at the Georgia Genomics & Bioinformatics Core (GGBC) on a NextSeq500 platform (Illumina). Average sequencing depth was ∼41.1x among samples (Table S7).

MethylC-seq reads were processed and aligned using Methylpy v1.3.2 [69]. Default parameters were used expect for the following: clonal reads were removed, lambda DNA was used as the unmethylated control, and binomial test was performed for all cytosines with at least three mapped reads.

### Identification of Differentially Methylated Regions

Identification of differentially methylated regions (DMRs) was performed using Methylpy v1.3.2 [69]. All methylome samples were analyzed together to conduct an undirected identification of DMRs across all samples in the CNN (N=A, C, G, T) context. Default parameters were used. Only DMRs at least 40-bp long with at least three differentially methylated cytosines (DMS) and five or more cytosines with at least one read were retained. For each DMR, the weighted methylation level was computed as mC / (mC + uC) where mC and uC are the number of reads supporting a methylated cytosine and unmethylated cytosine, respectively [41].

To identify epigenetic variants in these samples, we used a one-sided z-test to test for a significant difference in methylation level of DMRs pairwise between branches. For each pair, only DMRs with at least 5% difference in methylation level were used, regardless of underlying context. Resulting *P* values were adjusted using Benjamini-Hochberg correction (N = 383,600) with FDR = 0.05 [70] and DMRs are defined by adjusted *P* value ≤ 0.05.

### Identification of Methylated Regions

For each sample, an unmethylated methylome was generated by setting the number of methylated reads to zero while maintaining the total number of reads. Methylpy DMR identification program [69] was applied to each sample using the original methylome and unmethylated methylome with the same parameters as used for DMR identification. Regions less than 40 bp-long, fewer than three DMS, and fewer than five cytosines with at least one read were removed. Remaining regions from all samples were merged using BEDtools v2.27.1 [71].

### Assigning genomic features to DMRs

A genomic feature map was created such that each base pair of the genome was assigned a single feature type (transposable element/repeat, promoter, untranslated region, coding sequence, and intron) based on the previously described annotation. Promoters were defined as 2 kb upstream of the transcription start site of protein-coding genes. At positions where multiple feature types could be applicable, such as a transposon in an intron or promoter overlapping with adjacent gene, priority was given to untranslated regions (highest), introns, coding sequences, promoter, and transposon (lowest). Positions without an assignment were considered intergenic. Genomic feature content of each DMR and methylated region was assigned proportionally based on the number of bases in each category.

### Identification of pseudo-allele methylation

We aimed to categorize the DMRs into three pseudo-allele states: homozygous methylated, heterozygous, and homozygous unmethylated. First, DMRs were filtered on the following criteria: i) at least 25% change in weighted CG methylation level between the highest and lowest methylation level of the samples; ii) at least one sample had a CG methylation level of at least 75%; and iii) at least two “covered” CG positions. A “covered” CG is defined as having at least one read for both symmetrical cytosines in all samples. After filtering, 4,488 regions were used for analysis.

For each region in each sample, we next categorize the aligned reads overlapping the region. If at least 35% of its “covered” CG sites are methylated, the read is categorized as methylated. Otherwise it is an unmethylated read. Finally, we define the pseudo-allele state by the portion of methylated reads; homozygous unmethylated: ≤ 25%, heterozygous: > 25% and < 75%, and homozygous methylated: ≥ 75%.

The null distribution was created by randomly shuffling the filtered DMRs in the genome such that each simulated region is the same length as the original and it has at least two “covered” CGs. The above procedure was applied and number of epigenotype changes was determined. This was repeated for a total of 10 times.

The following special classes of DMRs were identified: highly variable, single loss, single gain, and tree specific. A DMR is highly variable if there were pseudo-allele changes between all adjacent branches. A DMR is single loss if all but one branch was homozygous methylated, and one was homozygous unmethylated. Similarly, a DMR is single gain if all but one branch was homozygous unmethylated and one branch was homozygous methylated. Finally, a DMR is “tree specific” if all tree 13 branches were homozygous unmethylated and all tree 14 branches were homozygous methylated or vice versa.

### Estimating somatic epimutation rate

We previously developed a method for estimating ‘germline’ epimutation rates in *A. thaliana* based on multi-generational methylation data from Mutation Accumulation lines [32]. In a companion method paper to the present study (Shahryary et al. 2019, co-submission), we have extended this approach to estimating somatic epimutation rates in long-lived perennials such as trees using leaf methylomes and coring data as input.

This new inference method, which we call *AlphaBeta*, treats the tree branching structure as a pedigree of somatic lineages using the fact that these cell lineages carry information about the epimutational history of each branch. *AlphaBeta* is implemented as a bioconductor R package (http://bioconductor.org/packages/devel/bioc/html/AlphaBeta.html). Using this approach, we estimate somatic epimutation rates for individual CG, CHG, and CHH sites independently, but also for regions. For the region-level analysis, we first use the differentially methylated regions (DMRs) identified above. Sampling from the distribution of DMR sizes, we then split the remainder of the genome into regions, which we refer to as “non-DMRs”. Per sample, we aggregate the total number of methylated Cs and unmethylated Cs in each region corresponding to a DMRs or a non-DMRs and used these counts as input for *AlphaBeta*.

### mRNA-seq sequencing and analysis

Total RNA was extracted from leaf tissue in each branch using the Direct-zol RNA MiniPrep Plus kit (Zymo Research) with Invitrogen’s Plant RNA Reagent. Total RNA quality and quantity were assessed before library construction. Strand-specific RNA-seq libraries were constructed using the TruSeq Stranded mRNA LT kit (Illumina) following the manufacturer’s instructions. For each sample, three independent libraries (technical replicates) were constructed. Libraries were sequenced to paired-end 75-bp reads at the GGBC on a NextSeq500 platform (Illumina). Summary statistics are included in the Table S9.

For analysis, first, paired-end reads were trimmed using Trimmomatic v0.36 [72]. Trimming included removing TruSeq3 adapters, bases with quality score less than 10, and any reads less than 50-bp long. Second, remaining reads were mapped to the *Stettler* genome with HiSAT2 [73] using default parameters except to report alignments for transcript assemblers (--dta). The HiSAT2 transcriptome index was created using extracted splice sites and exons from the gene annotation as recommended. Last, transcriptional abundances for genes in the reference annotation were computed for each sample using StringTie v1.3.4d [74]. Default parameters were used except to limit estimates to reference transcripts. TPM (transcripts per million) values were outputted to represent transcriptional abundance.

### Identification of differentially expressed genes

Differentially expressed genes (DEGs) were identified using DeSeq2 v1.22.2 [46]. The count matrix was extracted from StringTie output files and the analysis was performed using the protocol (ccb.jhu.edu/software/stringtie/index.shtml?t=manual#deseq). Abundances for all samples were joined into one DESeq dataset with α = 0.01. Gene abundance was compared between all samples pairwise. In each pair, a gene was considered differentially expressed if the adjusted *P* value ≤ 0.01 and the log_2_-fold change ≥ 1. Genes differentially expressed in any pair were included for subsequent analysis.

### Overlap of DMRs and DEGs

We identified DMRs which overlapped the promoter region (2 kb upstream of transcription start site) and gene body of annotated genes. For each DMR-gene pair, we computed the Pearson’s product moment correlation coefficient between the weighted methylation level of the DMR and average gene abundance among replicates in TPM. Next, looking only at genes which were previously identified as differently expressed, we performed a two-sided Pearson’s correlation test for each DMR-DEG pair to test for statistically significant correlations. Resulting *P* values were multiple test corrected with Benjamini-Hochberg correction (N = 382, FDR = 0.05) [70]. Adjusted *P* values ≤ 0.05 were considered significantly correlated.

## DECLARATIONS

### Ethics approval and consent to participate

Not applicable

### Consent for publication

Not applicable

### Availability of data and materials

Raw sequence data used for genome assembly, resequencing and identification of structural variation of individual branches are available at NCBI SRA (PRJNA516415). Raw sequence data for whole-genome bisulfite sequencing and mRNA-sequencing are available in GEO under accession GSE132939.

Custom analysis scripts used in this study are available in the GitHub repository https://github.com/schmitzlab/somatic-epigenetic-mutation-poplar.

### Authors’ Contributions

RJS, FJ, GAT, RS and JS conceived and designed the experiments. JG, SS, KB, KL, CA, AL, DK, JT, RW performed data generation. BTH, JD, MCT, YS, RH, SM, JJ, PPG, FJ performed data analysis. BTH prepared the figures and manuscript. BTH, DWH, GAT, FJ, and RJS wrote and revised the manuscript with input from all authors. All authors read and approved the final manuscript.

## Acknowledgements

We thank the JGI and collaborators for pre-publication access to *P. trichocarpa* v4.0 genome sequence for chromosome scale ordering of the *Stettler* genome and use of the RNA-seq data from the JGI Plant Gene Atlas for annotation. Sample collection was supported by the Center for Bioenergy Innovation (CBI). CBI is Bioenergy Research Centers supported by the Office of Biological and Environmental Research in the US Department of Energy Office of Science. We also thank Dr. Pankaj Jaiswal for use of additional RNA-seq data included in the annotation.

