## Supplementary Material for "The somatic genetic and epigenetic mutation rate in a wild long-lived perennial *Populus trichocarpa*"

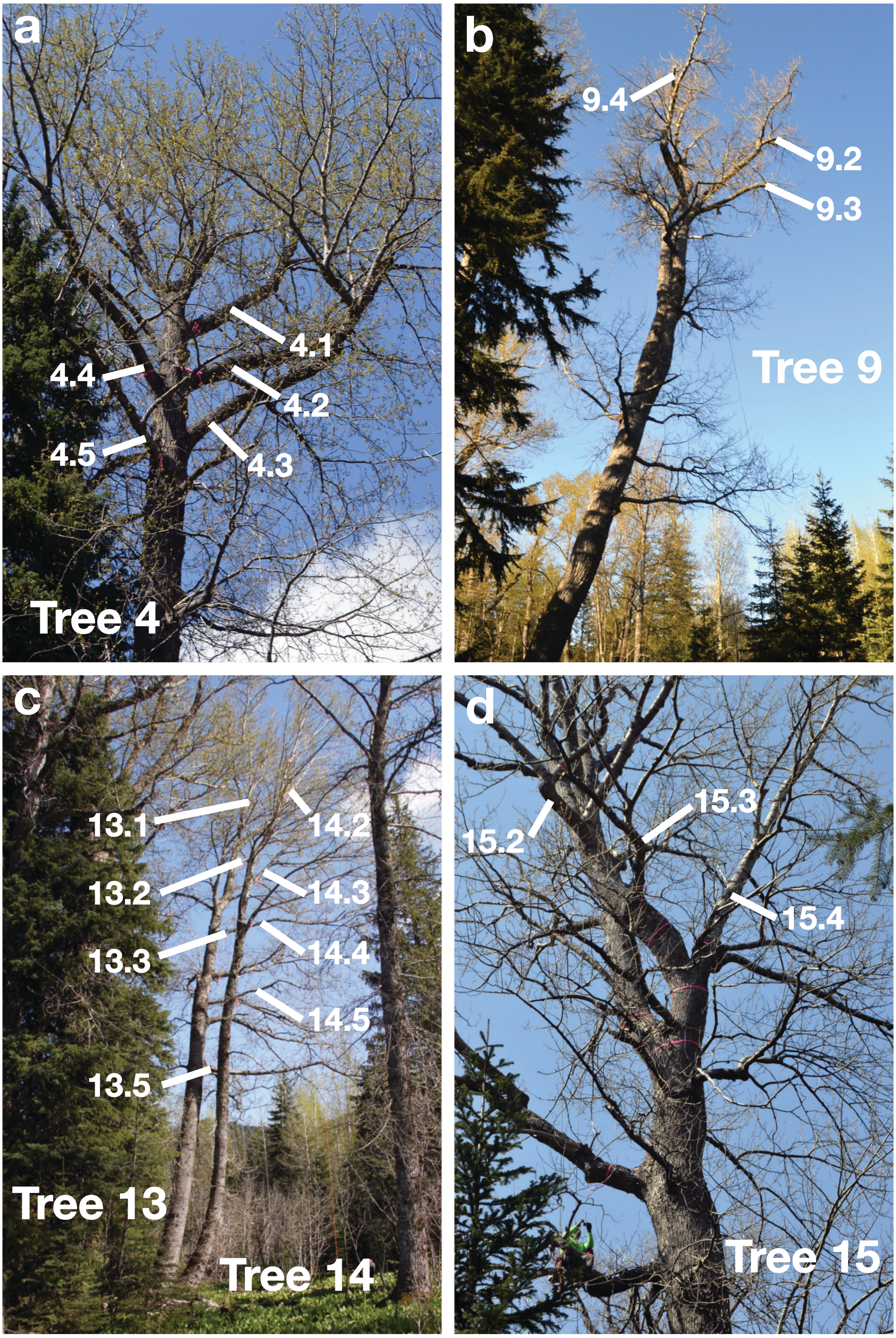


**Fig. S1. Photographs of the trees used in this study.** Photographs of tree 4 (a), tree 9 (b), tree 13 and 14 (c), and tree 15 (d) with branches labeled. Leaf samples were collected from each branch.

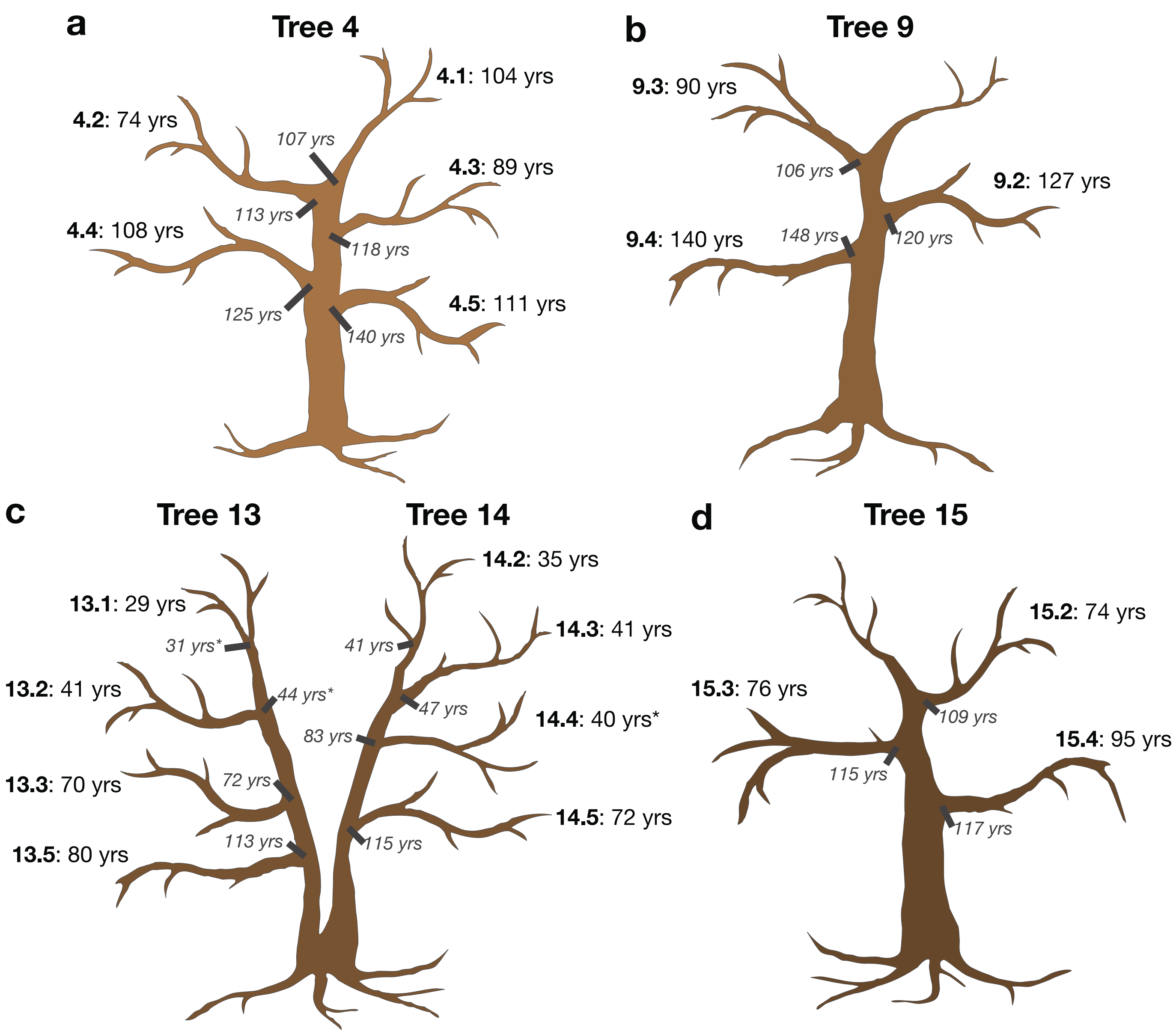


**Fig. S2. Schematic drawings of additional trees in the study.** Schematic drawings of tree 4 (a), tree 9 (b), tree 13 and 14 (c), and tree 15 (d) with estimated terminal branch ages and age where branch meets the main stem (gray italic). Leaf samples were collected from each branch for genomic sequencing libraries.

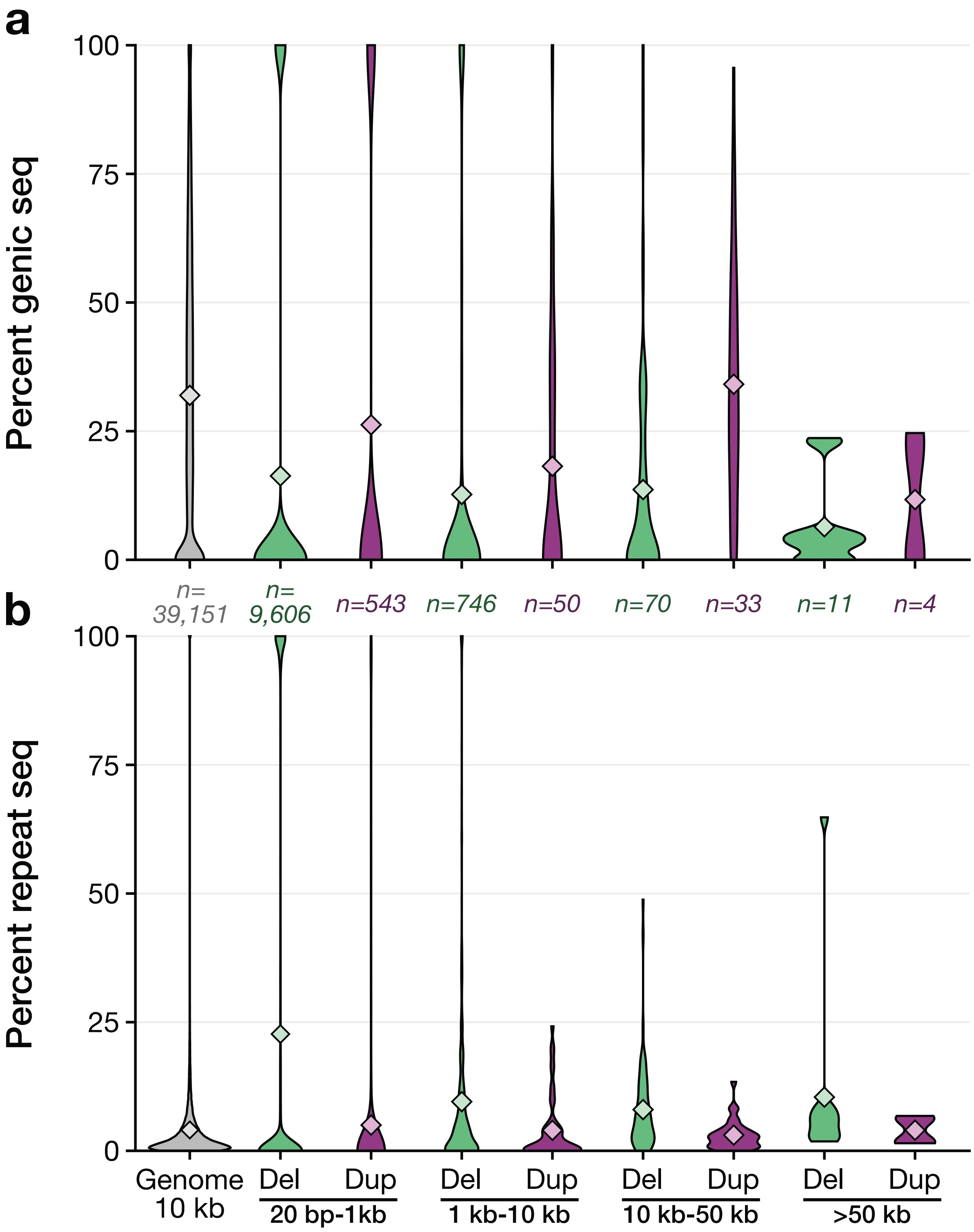


**Fig. S3. Duplications contain a higher proportion of genic sequences and deletions contain a higher proportion of repeat sequence.** a) For deletions (Del, green) and duplications (Dup, purple) structural variants grouped by size, distribution of the proportion of the SV sequence that overlaps with an annotate gene. Same as a except proportion of the SV sequence that overlaps transposons and repeat sequences. Genome-null (gray) is measured for 10-kb windows across the genome. Diamond represents the group mean. Number of SVs in each group is specified above b.

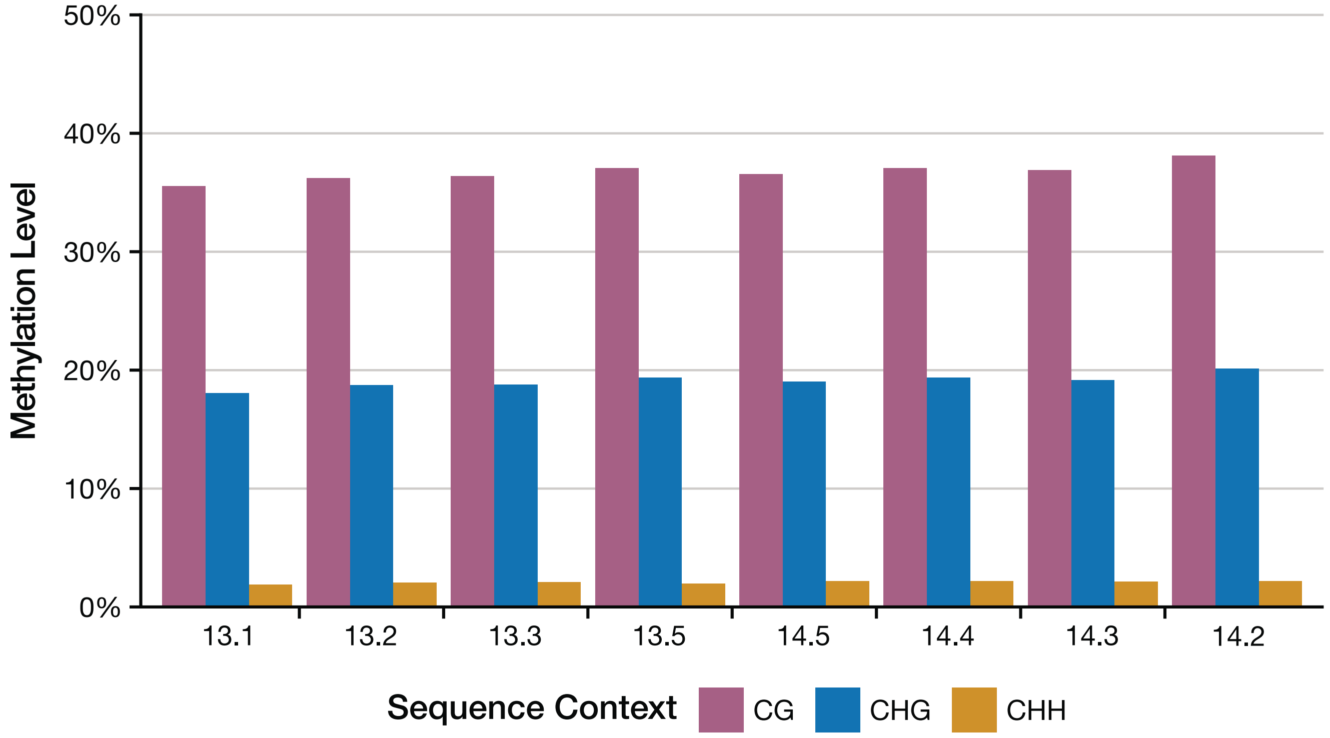


**Fig. S4. Genome weighted methylation levels.** Genome-wide weighted methylation level for mCG (red), mCHG (blue), and mCHH (yellow) for samples in tree 13 and tree 14.

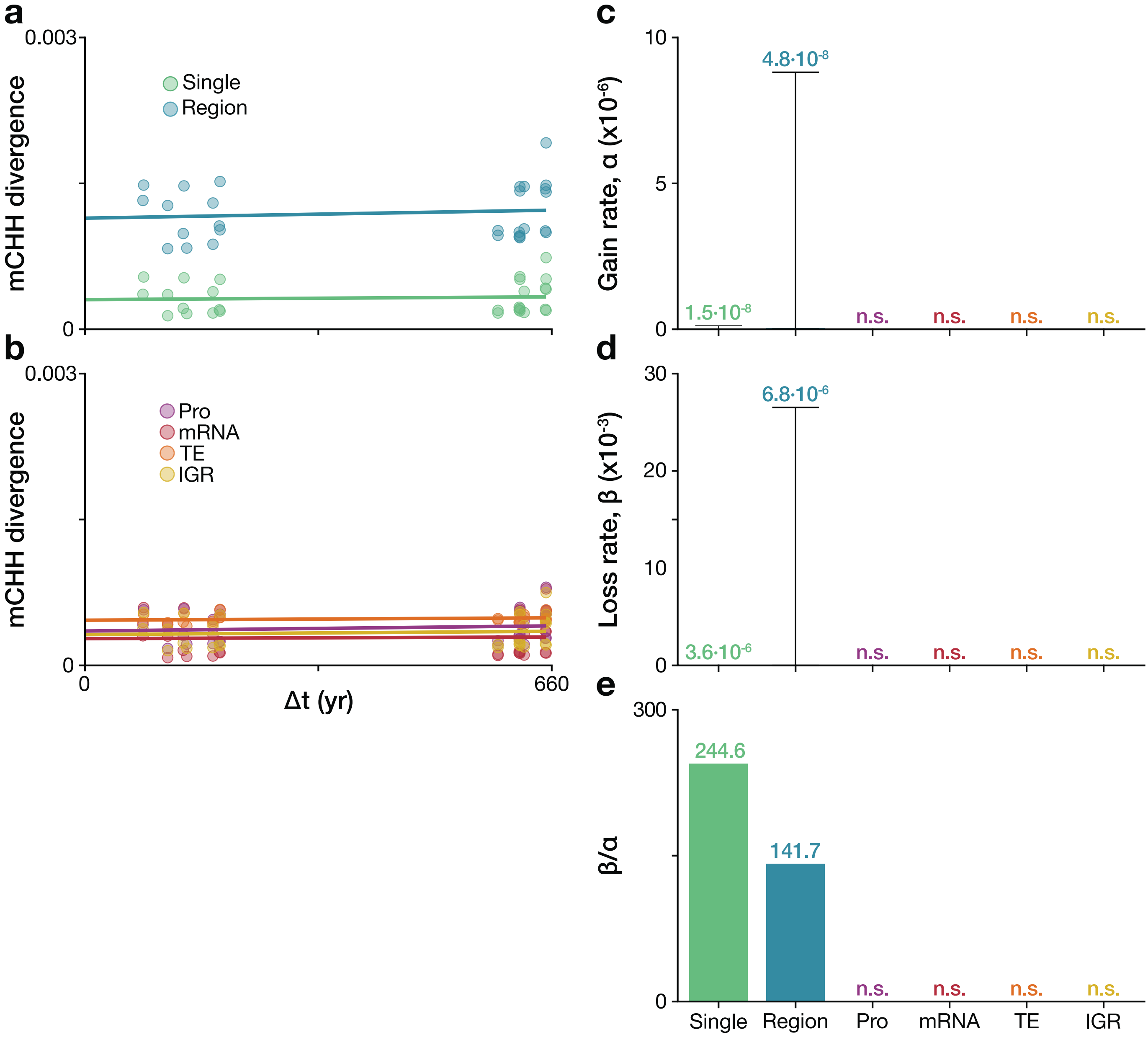


**Fig. S5. Somatic epimutation rates for single sites, regions, and by genomic feature in the CHH context.** Methylation divergence by branch time divergence for single sites and regions (a) and genomic features (b). Abbreviations: Pro, promoter; 1.5 kb upstream of TSS; TE, transposable elements and repeats; and IGR, intergenic regions. c) Estimated methylation gain rate, α, by feature. d) Estimated methylation loss rate, β, by feature. e) Estimated ratio of loss to gain, β/α. If there is no significant effect of branch age for the feature, it is marked n.s.

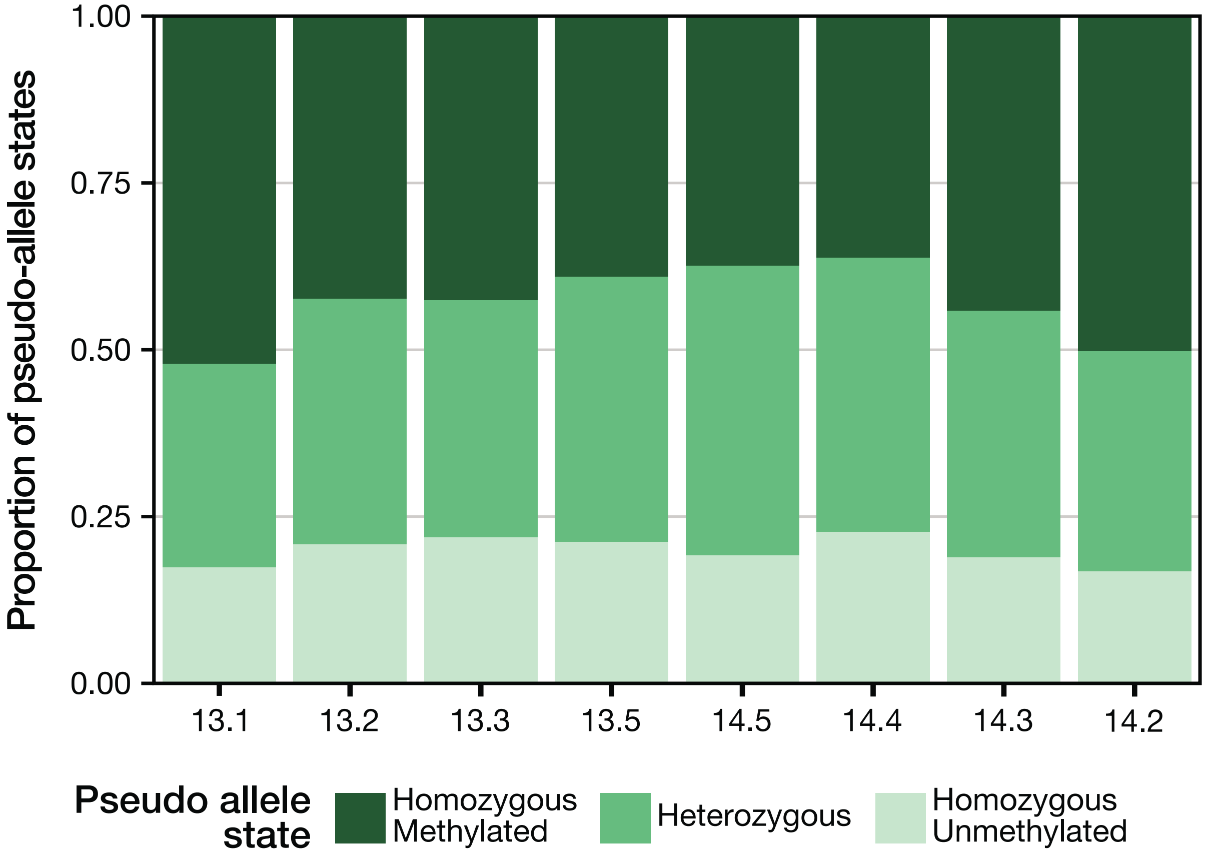


**Fig. S6. Pseudo allele states of DMRs among samples.** Branches 13.1 and 14.2 have more homozygous methylated pseudo alleles than the older branches. For each sample, the distribution of tested DMRs (N = 4,488) assignments. Possible pseudo allele states are homozygous methylated (dark green), heterozygous (medium green), and homozygous unmethylated (light green).

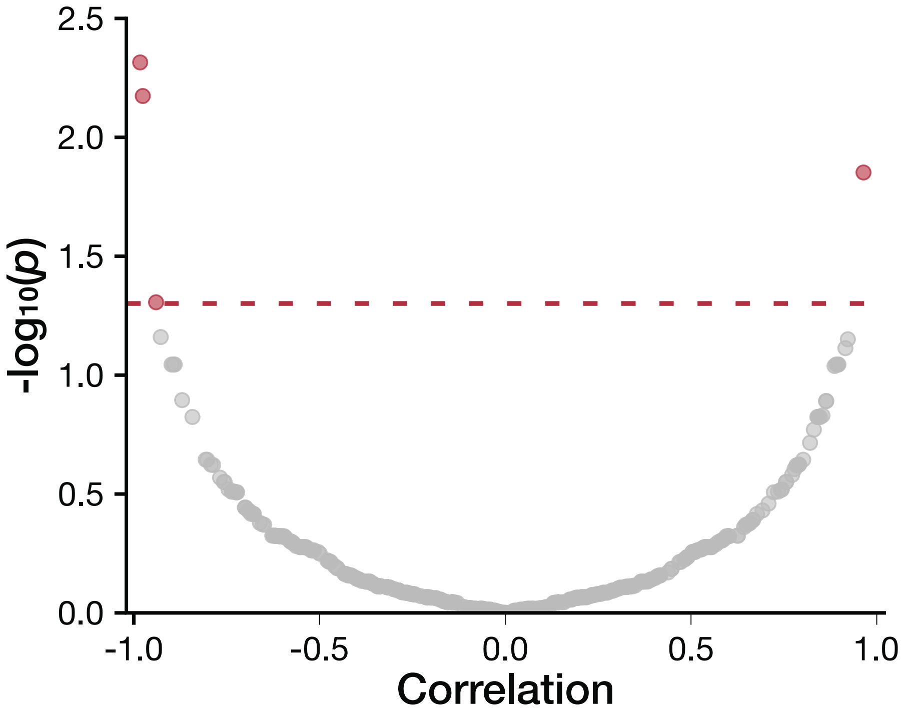


**Fig. S7. Gene expression of differentially expressed genes is rarely correlated to methylation level of nearby differentially methylated regions.** Each point represents a differentially expressed gene-differentially methylated region pair where the DMR is in the gene body or within 2-kb upstream. Correlation is the Pearson's correlation between gene expression, average of replicates as TPM, and weighted methylation level. Pearson's correlation test, two-sided, was performed on each pair then multiple test corrected using Benjamini-Hochberg (N = 382, FDR=0.05). Red dashed line is the significance threshold, adjusted *P* value ≤ 0.05. Significant DEG-DMR pairs are colored red.

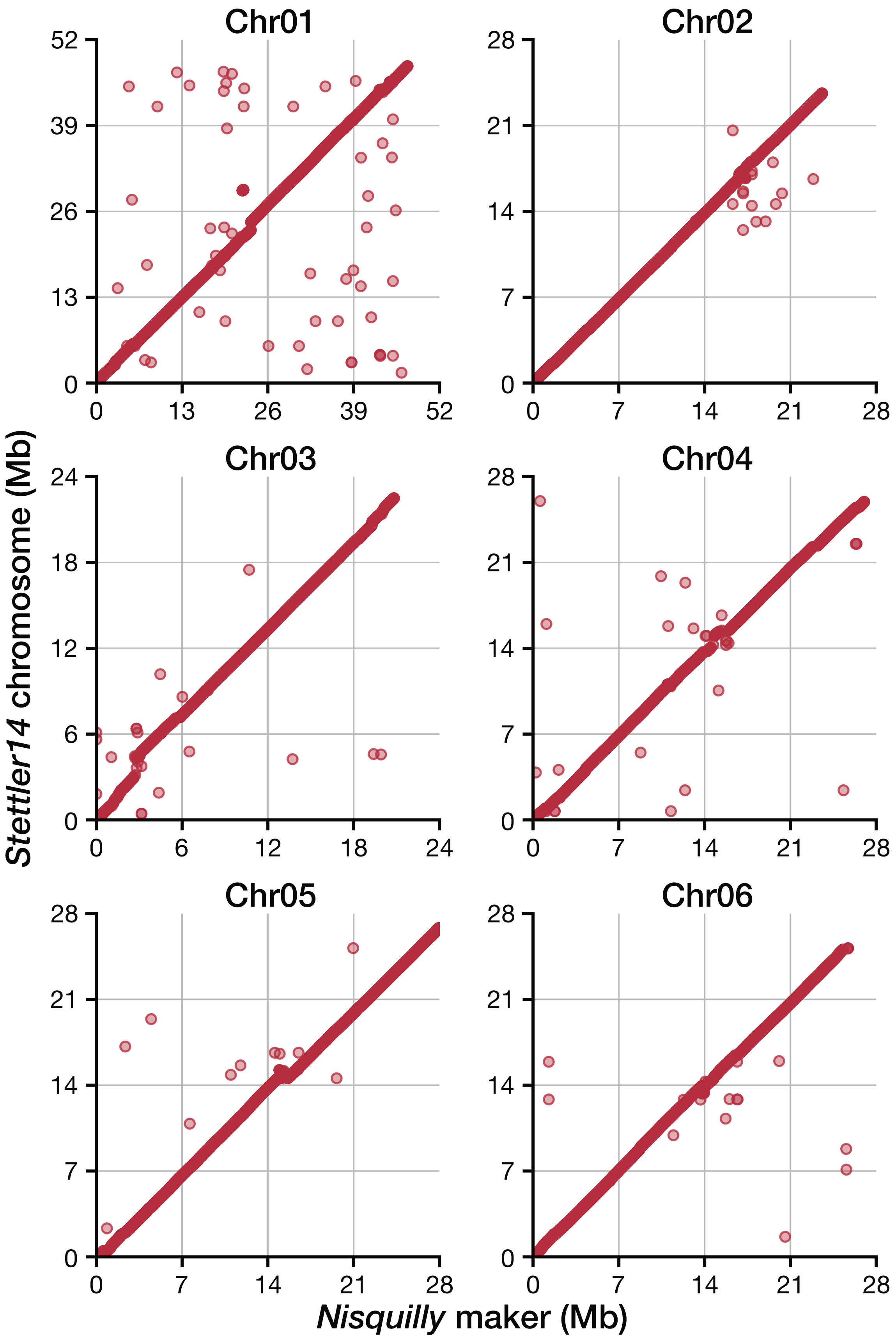


**Fig. S8.** Syntenic *Nisqually* marker placements on the *Populus trichocarpa* var. *Stettler* chromosomes. Each point represents a *Nisqually* marker positioned along the *Nisqually* chromosome along the x-axis and *Stettler* chromosome along the y-axis.

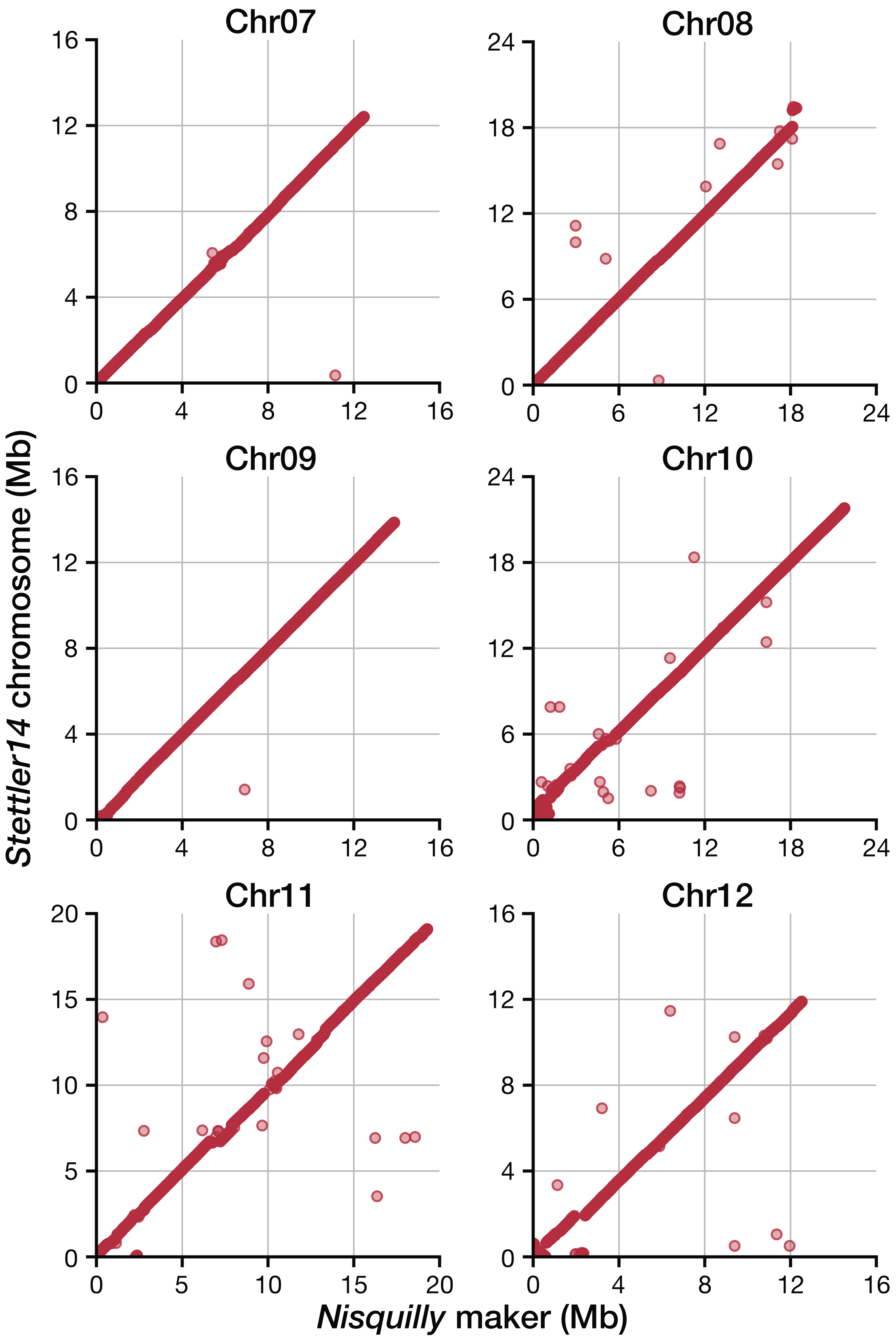


**Fig. S8.** Syntenic *Nisqually* marker placements on the *Populus trichocarpa* var. *Stettler* chromosomes. Each point represents a *Nisqually* marker positioned along the *Nisqually* chromosome along the x-axis and *Stettler* chromosome along the y-axis.

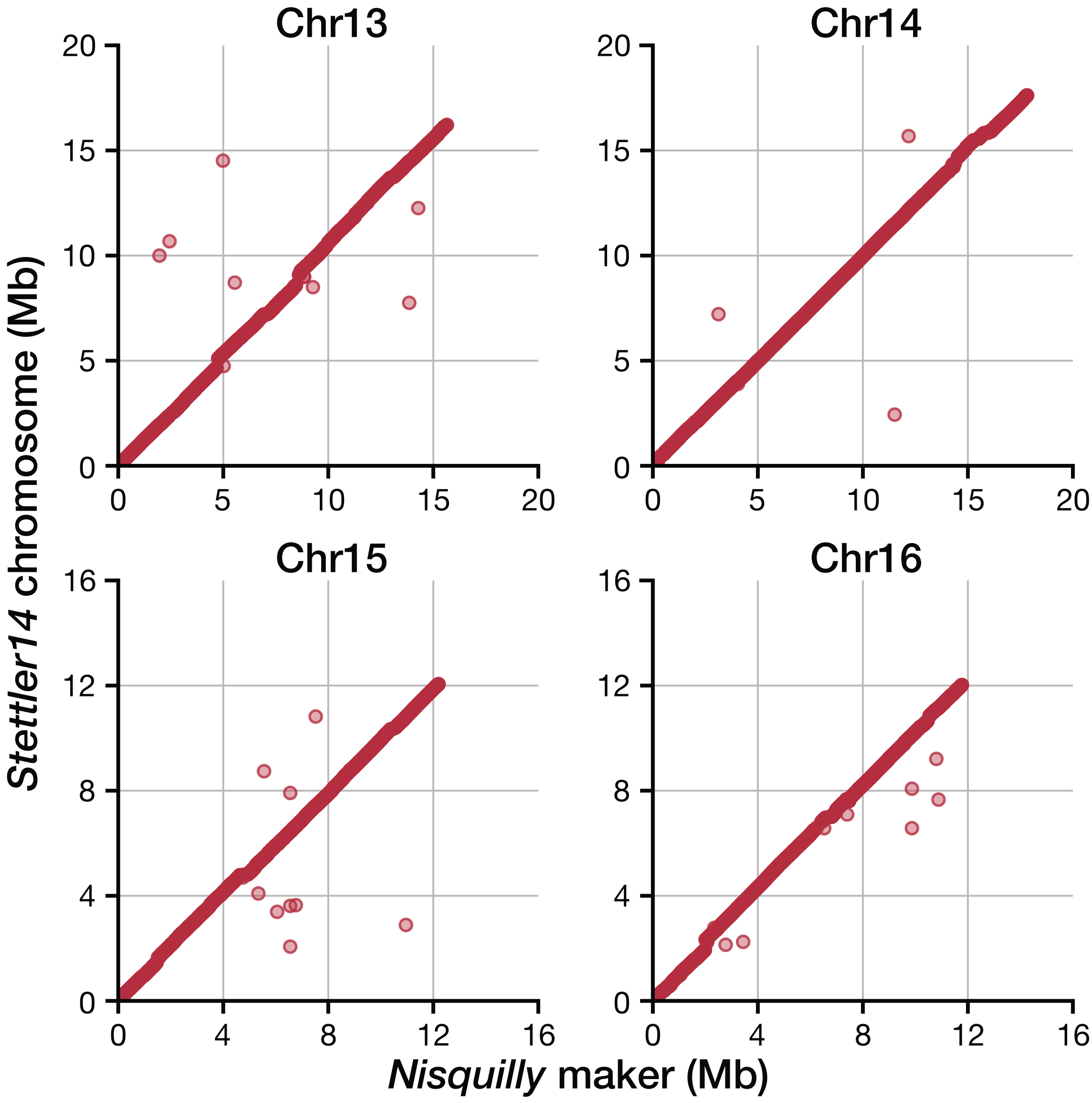


**Fig. S8.** Syntenic *Nisqually* marker placements on the *Populus trichocarpa* var. *Stettler* chromosomes. Each point represents a *Nisqually* marker positioned along the *Nisqually* chromosome along the x-axis and *Stettler* chromosome along the y-axis.

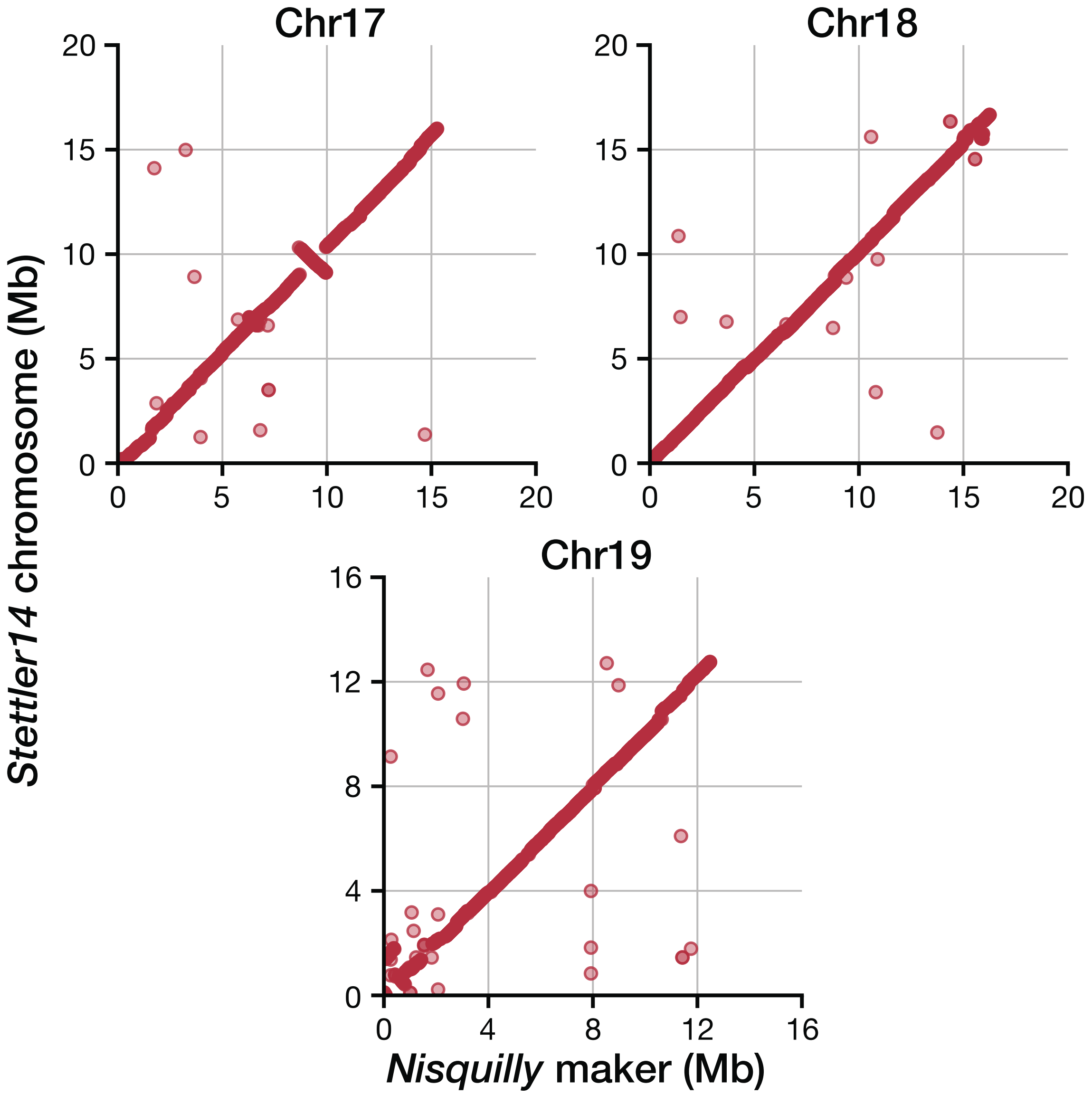


**Fig. S8.** Syntenic *Nisqually* marker placements on the *Populus trichocarpa* var. *Stettler* chromosomes. Each point represents a *Nisqually* marker positioned along the *Nisqually* chromosome along the x-axis and *Stettler* chromosome along the y-axis.

**Table S1.** High-confidence SNPS identified in Tree 13 with corresponding branch genotypes. In the genotypes, "RR" is homozygous for reference allele and "RA" is heterozygous for alternative allele.

| Chrm | Position | Ref | Alt | 13.1 Genotype | 13.2 Genotype | 13.3 Genotype | 13.5 Genotype | Genomic Feature | Feature Name |
|---|---|---|---|---|---|---|---|---|---|
| Chr01 | 3,680,749 | T | A | RR | RR | RR | RR | TE | Ogre-PT3_I-int |
| Chr01 | 13,084,368 | T | G | RR | RA | RR | RR | TE | Helitron-1_PTr |
| Chr01 | 17,000,961 | G | A | RR | RR | RR | RA | TE | Copia45-PTR_I-int |
| Chr01 | 18,843,033 | C | T | RR | RR | RR | RA | TE | Gypsy-39_PT-I |
| Chr01 | 19,584,181 | G | A | RR | RR | RR | RA | TE | Copia-3_PTri-I |
| Chr01 | 20,097,387 | C | T | RR | RA | RR | RR | TE | Gypsy-78_PTr-LTR |
| Chr01 | 20,099,693 | C | T | RA | RR | RR | RR | intergenic | |
| Chr01 | 21,841,528 | C | T | RR | RR | RR | RA | promoter | PtStettler14.01G189300 |
| Chr01 | 22,889,602 | T | G | RR | RA | RR | RR | TE | Gypsy7-PTR_I-int |
| Chr01 | 27,908,671 | C | T | RR | RA | RR | RR | intergenic | |
| Chr01 | 28,693,440 | C | T | RR | RA | RR | RR | TE | hAT-7_PTr |
| Chr01 | 29,156,750 | G | A | RR | RR | RA | RR | promoter | PtStettler14.01G238100 |
| Chr01 | 30,890,308 | T | C | RR | RR | RR | RR | TE | POPGY2_LTR |
| Chr01 | 34,853,944 | G | C | RR | RA | RR | RR | TE | EnSpm3_PT |
| Chr01 | 35,718,549 | A | T | RR | RR | RR | RA | intergenic | |
| Chr01 | 36,766,170 | C | T | RA | RA | RR | RR | TE | Helitron-1_PTr |
| Chr01 | 37,908,654 | G | A | RR | RR | RA | RR | TE | POPCOP2_I-int |
| Chr01 | 38,894,850 | T | G | RA | RR | RR | RR | intergenic | |
| Chr01 | 43,319,055 | G | A | RA | RR | RR | RR | TE | Ogre-PT1_I-int |
| Chr01 | 50,383,082 | C | T | RA | RA | RR | RR | intergenic | |
| Chr02 | 11,217,365 | T | C | RR | RA | RA | RR | TE | Copia29-PTR_I-int |
| Chr02 | 11,670,762 | C | T | RA | RR | RR | RR | intergenic | |
| Chr03 | 1,232,072 | A | G | RR | RA | RR | RR | TE | Copia-56_PTr-I |
| Chr03 | 2,727,138 | C | A | RR | RR | RR | RA | TE | EnSpm3_PT |

| | | | | | | | | | |
|---|---|---|---|---|---|---|---|---|---|
| Chr03 | 8,160,019 | G | T | RA | RR | RR | RR | intergenic | |
| Chr03 | 13,176,335 | C | T | RR | RA | RR | RR | TE | Helitron-N1_PTr |
| Chr03 | 15,811,182 | C | A | RA | RR | RR | RR | intergenic | |
| Chr03 | 20,942,498 | G | T | RR | RR | RR | RA | TE | SINE2-1_PTr |
| Chr04 | 2,007,485 | T | G | RA | RR | RR | RR | TE | Copia-56_PTr-I |
| Chr04 | 4,310,636 | T | A | RR | RR | RA | RR | mRNA | PtStettler14.04G046600 |
| Chr04 | 5,084,143 | G | A | RA | RR | RR | RR | intergenic | |
| Chr04 | 6,419,346 | A | T | RR | RA | RR | RR | mRNA | PtStettler14.04G067400 |
| Chr04 | 6,750,621 | G | A | RA | RR | RR | RR | intergenic | |
| Chr04 | 11,925,626 | G | A | RA | RR | RR | RR | TE | Gypsy-79_PTr-LTR |
| Chr04 | 12,602,059 | C | T | RR | RR | RR | RA | TE | ENSPM1_PT |
| Chr04 | 14,664,569 | G | A | RR | RR | RR | RR | TE | Gypsy21-PTR_I-int |
| Chr04 | 16,751,386 | G | A | RR | RR | RA | RR | intergenic | |
| Chr04 | 23,145,609 | A | G | RR | RA | RR | RR | TE | Gypsy-73_PTr-LTR |
| Chr05 | 724,815 | A | G | RR | RA | RR | RR | mRNA | PtStettler14.05G010800 |
| Chr05 | 2,530,404 | G | A | RA | RR | RR | RR | intergenic | |
| Chr05 | 4,349,460 | T | A | RR | RA | RR | RR | intergenic | |
| Chr05 | 4,841,986 | G | A | RR | RR | RR | RA | intergenic | |
| Chr05 | 6,750,399 | G | A | RR | RA | RR | RR | intergenic | |
| Chr05 | 7,678,555 | T | C | RA | RA | RR | RR | mRNA | PtStettler14.05G094400 |
| Chr05 | 9,053,607 | G | A | RR | RR | RR | RA | intergenic | |
| Chr05 | 10,930,336 | G | T | RR | RR | RA | RR | intergenic | |
| Chr05 | 16,304,960 | G | A | RR | RR | RR | RA | intergenic | |
| Chr05 | 22,750,874 | C | T | RR | RR | RR | RA | TE | DNA-3-1_PTr |
| Chr06 | 3,968,535 | G | A | RR | RR | RA | RR | promoter | DNA-3-1_PTr |
| Chr06 | 8,430,261 | A | G | RR | RR | RA | RR | TE | DNA-3-3_PTr |
| Chr06 | 9,721,447 | A | G | RR | RR | RA | RR | mRNA | PtStettler14.06G107700 |
| Chr06 | 13,034,017 | G | A | RR | RA | RR | RR | TE | Ogre-PT1_I-int |

| | | | | | | | | | |
|---|---|---|---|---|---|---|---|---|---|
| Chr06 | 13,156,719 | T | A | RR | RA | RR | RR | mRNA | PtStettler14.06G133600 |
| Chr06 | 21,337,989 | G | A | RA | RA | RA | RR | TE | Helitron-N1_PTr |
| Chr06 | 22,272,727 | G | A | RA | RA | RA | RR | intergenic | |
| Chr06 | 23,593,822 | G | A | RR | RA | RR | RR | mRNA | PtStettler14.06G206800 |
| Chr07 | 1,880,034 | G | A | RA | RR | RR | RR | promoter | PtStettler14.07G023100 |
| Chr07 | 2,168,501 | C | T | RR | RA | RR | RR | mRNA | PtStettler14.07G026600 |
| Chr07 | 5,569,304 | G | T | RR | RA | RR | RR | promoter | PtStettler14.07G049500 |
| Chr07 | 6,408,023 | G | A | RR | RR | RA | RR | TE | Gypsy-25_PTr-I |
| Chr07 | 12,690,085 | A | C | RR | RA | RR | RR | mRNA | PtStettler14.07G097000 |
| Chr08 | 4,851,487 | C | T | RA | RR | RR | RR | TE | DNA-3-2B_PTr |
| Chr08 | 15,270,575 | T | A | RR | RA | RR | RR | intergenic | |
| Chr08 | 16,874,471 | T | C | RR | RA | RR | RR | TE | Gypsy-79_PTr-LTR |
| Chr08 | 16,938,813 | C | T | RA | RR | RR | RR | intergenic | |
| Chr08 | 17,164,868 | G | A | RA | RR | RR | RR | intergenic | |
| Chr10 | 223,598 | G | T | RR | RR | RA | RR | intergenic | |
| Chr10 | 2,645,988 | T | C | RR | RA | RR | RR | intergenic | |
| Chr10 | 5,400,307 | C | T | RA | RA | RR | RR | intergenic | |
| Chr10 | 5,638,948 | C | T | RR | RR | RR | RA | TE | Gypsy18-PTR_I-int |
| Chr10 | 8,199,692 | C | T | RR | RA | RR | RR | promoter | PtStettler14.10G041500 |
| Chr10 | 18,253,568 | G | T | RA | RR | RR | RR | intergenic | |
| Chr11 | 3,067,101 | A | C | RR | RR | RA | RR | intergenic | |
| Chr11 | 6,879,079 | C | A | RR | RA | RR | RR | mRNA | PtStettler14.11G060200 |
| Chr11 | 11,521,364 | G | A | RR | RA | RR | RR | intergenic | |
| Chr11 | 12,995,509 | A | C | RR | RR | RR | RR | TE | Gypsy-30_PTr-LTR |
| Chr11 | 15,878,255 | T | C | RR | RR | RR | RR | TE | Gypsy-71_PTr-LTR |
| Chr11 | 18,064,447 | T | C | RA | RA | RA | RR | intergenic | |
| Chr12 | 3,178,343 | C | T | RR | RR | RR | RA | TE | Gypsy-26_PTr-LTR |
| Chr12 | 7,107,969 | T | C | RR | RR | RA | RR | intergenic | |

| | | | | | | | | | |
|---|---|---|---|---|---|---|---|---|---|
| Chr12 | 10,764,498 | G | A | RR | RR | RA | RR | TE | Helitron-N3_PTr |
| Chr12 | 12,182,117 | A | G | RA | RR | RR | RR | intergenic | |
| Chr12 | 12,764,745 | T | A | RR | RR | RR | RR | TE | Copia43-PTR_I-int |
| Chr13 | 1,831,348 | G | A | RA | RR | RR | RR | intergenic | |
| Chr14 | 4,670,153 | A | G | RR | RA | RR | RR | TE | Copia13-PTR_I-int |
| Chr14 | 9,051,852 | C | T | RR | RA | RR | RR | intergenic | |
| Chr14 | 13,911,806 | A | C | RR | RR | RR | RA | TE | POPGY1_I-int |
| Chr14 | 15,926,176 | G | A | RR | RR | RR | RA | TE | Helitron-N1_PTr |
| Chr15 | 4,046,877 | A | G | RR | RR | RR | RR | intergenic | |
| Chr15 | 5,724,901 | A | T | RR | RR | RA | RR | intergenic | |
| Chr15 | 6,640,604 | C | T | RR | RA | RR | RR | TE | Gypsy-29_PTr-I |
| Chr15 | 7,447,713 | G | A | RR | RR | RR | RA | TE | Copia-93_PTr-LTR |
| Chr15 | 8,189,459 | C | T | RR | RA | RR | RR | mRNA | PtStettler14.15G051600 |
| Chr15 | 9,103,615 | G | A | RR | RR | RR | RA | promoter | PtStettler14.15G057400 |
| Chr15 | 9,280,803 | C | A | RA | RR | RR | RR | TE | EnSpm1B_PT |
| Chr15 | 11,916,633 | C | G | RR | RR | RA | RR | intergenic | |
| Chr15 | 12,184,591 | G | A | RR | RR | RR | RA | intergenic | |
| Chr15 | 12,599,915 | A | T | RR | RR | RR | RA | mRNA | PtStettler14.15G095000 |
| Chr16 | 4,655,795 | T | G | RR | RR | RR | RA | mRNA | PtStettler14.16G057300 |
| Chr16 | 5,850,725 | T | G | RR | RR | RR | RA | TE | Gypsy-76_PTr-LTR |
| Chr16 | 7,736,456 | C | T | RR | RA | RR | RR | promoter | PtStettler14.16G082600 |
| Chr16 | 7,781,247 | C | T | RA | RR | RR | RR | intergenic | |
| Chr16 | 8,036,782 | C | T | RR | RA | RR | RR | TE | Gypsy-74_PTr-I |
| Chr16 | 10,760,553 | G | A | RR | RR | RR | RA | TE | Helitron-N3_PTr |
| Chr17 | 5,464,790 | G | A | RR | RA | RR | RR | TE | Gypsy18-PTR_I-int |
| Chr17 | 7,196,478 | G | A | RR | RR | RA | RR | intergenic | |
| Chr17 | 7,446,065 | G | T | RR | RR | RR | RA | intergenic | |
| Chr17 | 10,671,769 | G | T | RA | RR | RR | RR | promoter | PtStettler14.17G074500 |

| | | | | | | | | | |
|---|---|---|---|---|---|---|---|---|---|
| Chr17 | 14,433,140 | G | A | RR | RA | RR | RR | intergenic | |
| Chr18 | 550,457 | C | A | RR | RR | RA | RR | mRNA | PtStettler14.18G005600 |
| Chr18 | 9,602,839 | T | A | RR | RR | RA | RR | TE | EnSpm3_PT |
| Chr18 | 13,302,911 | T | C | RR | RR | RR | RA | promoter | PtStettler14.18G096800 |
| Chr18 | 14,520,496 | G | T | RR | RA | RR | RR | mRNA | PtStettler14.18G111600 |
| Chr19 | 159,694 | G | A | RR | RR | RA | RR | TE | Gypsy-26_PTr-LTR |
| Chr19 | 895,499 | A | G | RR | RR | RA | RR | intergenic | |
| Chr19 | 10,872,272 | C | T | RA | RR | RR | RR | promoter | PtStettler14.19G065800 |
| Chr19 | 12,032,059 | C | A | RR | RR | RA | RR | intergenic | |
| Chr19 | 14,437,447 | G | A | RR | RR | RA | RR | TE | Copia46-PTR_I-int |

**Table S2.** High-confidence SNPS identified in Tree 14 with corresponding branch genotypes. In the genotypes, "RR" is homozygous for reference allele and "RA" is heterozygous for alternative allele.

| Chrm | Position | Ref | Alt | 14.5 Genotype | 14.4 Genotype | 14.3 Genotype | 14.2 Genotype | Genomic Feature | Feature Name |
|---|---|---|---|---|---|---|---|---|---|
| Chr01 | 3,021,991 | G | A | RR | RR | RA | RR | TE | Copia-92_PTr-I |
| Chr01 | 3,875,530 | G | A | RA | RR | RR | RR | TE | Copia-3_PTri-I |
| Chr01 | 6,123,545 | C | T | RA | RR | RR | RR | promoter | PtStettler14.01G068000 |
| Chr01 | 18,038,562 | T | G | RA | RR | RR | RR | TE | Gypsy18-PTR_I-int |
| Chr01 | 19,273,170 | C | T | RR | RA | RR | RR | TE | Gypsy-30_PTr-LTR |
| Chr01 | 19,500,304 | T | A | RR | RA | RR | RR | TE | Gypsy-25_PTr-LTR |
| Chr01 | 20,768,036 | A | T | RR | RR | RA | RR | TE | EnSpm3_PT |
| Chr01 | 21,180,079 | C | T | RR | RR | RA | RR | intergenic | |
| Chr01 | 25,248,811 | T | C | RR | RR | RR | RA | TE | POPGY1_I-int |
| Chr01 | 32,964,401 | C | T | RR | RA | RR | RR | TE | Copia-56_PTr-I |
| Chr01 | 34,106,911 | A | T | RR | RR | RR | RA | intergenic | |
| Chr01 | 35,795,970 | C | A | RR | RR | RA | RA | TE | Helitron-N1_PTr |
| Chr01 | 36,298,281 | C | T | RA | RR | RR | RA | TE | Copia9-PTR_I-int |

| | | | | | | | | | |
|---|---|---|---|---|---|---|---|---|---|
| Chr01 | 38,971,251 | T | A | RR | RR | RR | RR | TE | Gypsy18-PTR_I-int |
| Chr01 | 40,268,412 | G | A | RR | RA | RR | RR | intergenic | |
| Chr01 | 40,571,498 | G | T | RR | RA | RR | RR | mRNA | PtStettler14.01G332400 |
| Chr01 | 42,088,313 | T | A | RR | RR | RR | RA | intergenic | |
| Chr01 | 43,984,711 | T | C | RR | RR | RR | RR | promoter | PtStettler14.01G353800 |
| Chr01 | 44,305,840 | G | A | RR | RR | RR | RA | intergenic | |
| Chr02 | 489,916 | A | G | RR | RA | RR | RR | TE | Helitron-N3_PTr |
| Chr02 | 4,265,853 | C | T | RR | RR | RR | RA | promoter | hAT-1N_PTr |
| Chr02 | 5,336,055 | A | G | RR | RR | RA | RR | intergenic | |
| Chr02 | 5,371,222 | C | T | RR | RR | RR | RA | promoter | PtStettler14.02G071500 |
| Chr02 | 10,222,845 | T | C | RR | RR | RA | RR | TE | POPCOP1_I-int |
| Chr02 | 12,494,877 | G | A | RR | RR | RR | RA | TE | Helitron-1_PTr |
| Chr02 | 16,776,669 | C | T | RR | RR | RA | RR | promoter | PtStettler14.02G186200 |
| Chr02 | 18,652,325 | C | T | RR | RR | RA | RA | TE | Gypsy-27_PTr-LTR |
| Chr02 | 21,491,807 | T | C | RR | RR | RR | RA | TE | Gypsy-30_PTr-LTR |
| Chr02 | 21,672,642 | C | G | RR | RR | RA | RR | intergenic | |
| Chr02 | 25,291,486 | G | A | RR | RR | RA | RA | TE | Gypsy-39_PT-I |
| Chr03 | 325,130 | C | T | RR | RR | RA | RR | intergenic | |
| Chr03 | 3,823,775 | C | T | RR | RR | RR | RA | TE | Gypsy22-PTR_I-int |
| Chr03 | 7,342,298 | C | T | RR | RR | RA | RR | TE | SINE2-2_PTr |
| Chr03 | 9,438,180 | C | T | RR | RA | RR | RR | TE | Helitron-N3_PTr |
| Chr03 | 12,401,934 | A | G | RA | RR | RR | RR | mRNA | PtStettler14.03G073200 |
| Chr03 | 12,605,761 | A | G | RR | RA | RA | RA | intergenic | |
| Chr03 | 14,497,462 | G | C | RR | RR | RA | RR | mRNA | PtStettler14.03G096200 |
| Chr03 | 15,547,882 | G | A | RR | RA | RR | RR | intergenic | |
| Chr03 | 22,340,624 | T | G | RR | RR | RA | RR | intergenic | |
| Chr04 | 1,369 | T | A | RR | RA | RR | RR | intergenic | |
| Chr04 | 38,729 | G | A | RR | RA | RA | RA | TE | Gypsy-26_PTr-LTR |

| | | | | | | | | | |
|---|---|---|---|---|---|---|---|---|---|
| Chr04 | 778,883 | C | T | RR | RA | RR | RR | TE | Ogre-PT1_I-int |
| Chr04 | 2,651,778 | G | A | RR | RR | RR | RA | promoter | PtStettler14.04G030600 |
| Chr04 | 6,894,959 | G | C | RR | RA | RR | RR | TE | EnSpm1B_PT |
| Chr04 | 8,066,586 | G | A | RR | RA | RA | RR | intergenic | |
| Chr04 | 11,844,929 | G | A | RR | RR | RR | RA | TE | Gypsy-28_PTr-I |
| Chr04 | 12,667,744 | C | T | RR | RA | RR | RR | TE | Gypsy-27_PTr-I |
| Chr04 | 13,298,886 | C | A | RR | RR | RA | RR | intergenic | |
| Chr04 | 13,368,307 | T | C | RR | RA | RR | RR | TE | Gypsy-78_PTr-I |
| Chr04 | 13,756,856 | C | G | RR | RA | RR | RR | intergenic | |
| Chr04 | 17,448,141 | G | A | RR | RR | RA | RR | intergenic | |
| Chr04 | 19,020,051 | C | T | RR | RR | RR | RA | promoter | PtStettler14.04G154900 |
| Chr04 | 19,847,822 | C | G | RR | RR | RR | RR | mRNA | PtStettler14.04G164900 |
| Chr05 | 477,145 | C | T | RR | RA | RR | RR | TE | Copia46-PTR_I-int |
| Chr05 | 1,566,647 | G | A | RR | RR | RR | RA | TE | Gypsy-71_PTr-LTR |
| Chr05 | 1,655,537 | A | T | RR | RA | RR | RR | mRNA | PtStettler14.05G022700 |
| Chr05 | 5,104,250 | C | A | RR | RA | RR | RR | TE | Copia-93_PTr-LTR |
| Chr05 | 5,840,431 | T | C | RR | RR | RA | RR | intergenic | |
| Chr05 | 7,945,329 | C | T | RR | RA | RR | RR | TE | Copia-3_PTri-I |
| Chr05 | 8,553,289 | G | A | RR | RR | RA | RR | TE | Gypsy-75_PTr-LTR |
| Chr05 | 10,800,644 | G | A | RR | RR | RR | RA | TE | Caulimovirus-2_PTr |
| Chr05 | 13,876,857 | G | A | RR | RA | RR | RR | TE | Gypsy-35_PT-I |
| Chr05 | 17,269,379 | C | T | RR | RA | RR | RR | TE | hAT-5_PTr |
| Chr05 | 17,961,503 | A | G | RR | RR | RR | RR | intergenic | |
| Chr05 | 18,037,876 | T | G | RR | RR | RR | RA | intergenic | |
| Chr05 | 21,319,693 | T | A | RR | RA | RR | RR | mRNA | PtStettler14.05G187800 |
| Chr05 | 23,808,853 | C | G | RR | RA | RR | RR | intergenic | |
| Chr06 | 489,882 | C | T | RR | RA | RR | RR | TE | Gypsy-26_PTr-LTR |
| Chr06 | 6,649,018 | A | G | RR | RR | RR | RA | TE | Ogre-PT3_LTR |

| | | | | | | | | | |
|---|---|---|---|---|---|---|---|---|---|
| Chr06 | 8,432,769 | G | A | RR | RR | RA | RR | TE | DNA-3-1_PTr |
| Chr06 | 12,879,227 | C | T | RR | RA | RR | RR | intergenic | |
| Chr06 | 13,078,823 | C | T | RR | RR | RA | RR | promoter | PtStettler14.06G133200 |
| Chr06 | 13,113,622 | G | A | RR | RA | RR | RR | TE | ENSPM1_PT |
| Chr06 | 18,320,490 | C | T | RR | RR | RR | RA | intergenic | |
| Chr06 | 19,961,041 | T | A | RR | RA | RR | RR | intergenic | |
| Chr06 | 23,464,524 | C | T | RR | RR | RA | RR | intergenic | |
| Chr06 | 26,643,547 | C | T | RR | RA | RR | RR | intergenic | |
| Chr07 | 1,179,431 | T | C | RA | RR | RR | RR | mRNA | PtStettler14.07G014700 |
| Chr07 | 3,364,242 | G | T | RR | RR | RA | RR | intergenic | |
| Chr07 | 3,916,805 | G | A | RR | RR | RA | RA | promoter | PtStettler14.07G041300 |
| Chr07 | 3,954,801 | C | G | RR | RR | RA | RR | intergenic | |
| Chr07 | 5,211,186 | C | T | RR | RR | RA | RA | intergenic | |
| Chr07 | 6,021,662 | C | T | RR | RR | RR | RA | intergenic | |
| Chr07 | 8,131,543 | A | T | RR | RR | RR | RA | intergenic | |
| Chr07 | 8,689,627 | C | A | RR | RA | RR | RR | TE | EnSpm1B_PT |
| Chr07 | 10,999,644 | C | T | RR | RR | RR | RR | TE | Gypsy23-PTR_I-int |
| Chr07 | 14,430,205 | C | T | RR | RR | RA | RR | TE | Helitron-N2_PTr |
| Chr08 | 7,991,443 | G | A | RR | RA | RR | RR | promoter | PtStettler14.08G111000 |
| Chr08 | 9,172,616 | T | C | RR | RR | RR | RR | TE | Gypsy23-PTR_I-int |
| Chr08 | 11,966,249 | G | A | RR | RA | RR | RR | intergenic | |
| Chr08 | 14,099,204 | A | T | RR | RR | RR | RA | TE | EnSpm1B_PT |
| Chr08 | 17,740,276 | G | C | RR | RR | RR | RA | TE | hAT-3_PTr |
| Chr08 | 18,026,447 | G | T | RR | RA | RR | RR | TE | Caulimovirus-1_PTr |
| Chr08 | 18,830,197 | G | A | RR | RA | RR | RR | TE | EnSpm2_PTr |
| Chr08 | 18,830,272 | C | T | RR | RA | RR | RR | TE | EnSpm2_PTr |
| Chr08 | 19,613,054 | C | A | RR | RA | RR | RR | intergenic | |
| Chr09 | 577,123 | A | T | RR | RR | RR | RR | TE | Copia38-PTR_I-int |

| | | | | | | | | | |
|---|---|---|---|---|---|---|---|---|---|
| Chr09 | 2,321,924 | G | A | RR | RR | RR | RA | promoter | PtStettler14.09G009600 |
| Chr09 | 11,302,511 | T | C | RR | RR | RR | RA | TE | Copia19-PTR_I-int |
| Chr10 | 1,879,522 | G | T | RR | RR | RA | RR | TE | Gypsy-78_PTr-I |
| Chr11 | 2,434,004 | A | G | RR | RR | RR | RA | TE | Ogre-PT2_I-int |
| Chr11 | 2,588,249 | G | T | RA | RR | RR | RR | promoter | PtStettler14.11G025800 |
| Chr11 | 8,034,697 | C | T | RR | RA | RR | RR | TE | Helitron-1_PTr |
| Chr11 | 9,662,743 | A | G | RR | RA | RR | RR | TE | Gypsy-29_PTr-I |
| Chr11 | 12,727,094 | C | A | RR | RR | RA | RR | TE | Gypsy18-PTR_I-int |
| Chr11 | 14,264,394 | C | T | RR | RR | RA | RR | intergenic | |
| Chr11 | 16,481,007 | A | G | RR | RR | RR | RR | mRNA | PtStettler14.11G116100 |
| Chr12 | 94,956 | T | G | RR | RR | RA | RR | mRNA | PtStettler14.12G000800 |
| Chr12 | 192,041 | C | T | RR | RR | RR | RR | intergenic | |
| Chr12 | 4,325,795 | T | C | RR | RA | RR | RR | intergenic | |
| Chr12 | 7,146,575 | G | A | RR | RR | RA | RA | TE | ENSPM1_PT |
| Chr12 | 10,428,527 | C | T | RR | RR | RA | RA | promoter | PtStettler14.12G067400 |
| Chr13 | 1,689,009 | G | A | RR | RA | RR | RR | TE | DNA-3-2_PTr |
| Chr13 | 5,555,523 | G | A | RA | RR | RR | RR | TE | ENSPM1_PT |
| Chr13 | 7,123,591 | G | A | RR | RA | RR | RR | intergenic | |
| Chr13 | 7,127,671 | C | T | RR | RR | RA | RR | intergenic | |
| Chr13 | 8,610,319 | A | C | RR | RR | RR | RA | intergenic | |
| Chr13 | 10,412,391 | T | C | RR | RA | RA | RA | intergenic | |
| Chr13 | 13,734,311 | G | A | RA | RR | RR | RR | intergenic | |
| Chr13 | 15,498,389 | A | C | RR | RR | RA | RR | TE | Helitron-N4_PTr |
| Chr14 | 4,981,681 | G | A | RR | RR | RA | RA | TE | Helitron-1_PTr |
| Chr14 | 8,344,067 | G | A | RA | RR | RR | RR | TE | SINE2-2_PTr |
| Chr14 | 9,659,457 | C | T | RR | RR | RR | RA | intergenic | |
| Chr14 | 12,782,248 | G | A | RA | RR | RR | RR | intergenic | |
| Chr14 | 15,791,265 | C | T | RR | RR | RA | RR | intergenic | |

| | | | | | | | | | |
|---|---|---|---|---|---|---|---|---|---|
| Chr14 | 17,465,144 | A | G | RR | RA | RR | RR | TE | Ogre-PT3_I-int |
| Chr15 | 321,961 | G | A | RR | RA | RR | RR | promoter | PtStettler14.15G004400 |
| Chr15 | 4,557,354 | T | C | RR | RR | RR | RA | TE | Gypsy-79_PTr-LTR |
| Chr15 | 5,121,942 | G | A | RR | RR | RA | RR | TE | Copia14-PTR_I-int |
| Chr15 | 10,593,027 | A | C | RR | RA | RR | RR | TE | Copia38-PTR_I-int |
| Chr16 | 12,796,344 | G | A | RR | RR | RR | RA | TE | SINE2-2_PTr |
| Chr17 | 419,018 | A | G | RR | RR | RR | RA | TE | Copia-54_PTr-I |
| Chr17 | 3,268,172 | C | T | RR | RA | RR | RA | TE | Helitron-N1_PTr |
| Chr17 | 3,979,492 | C | T | RR | RR | RA | RR | TE | ENSPM1_PT |
| Chr17 | 4,938,558 | G | A | RA | RR | RR | RR | promoter | PtStettler14.17G046400 |
| Chr17 | 9,533,134 | C | T | RR | RA | RR | RR | intergenic | |
| Chr17 | 11,369,541 | G | A | RR | RA | RR | RR | TE | ENSPM1_PT |
| Chr19 | 982,282 | C | T | RR | RA | RR | RR | TE | Ogre-PT1_LTR |
| Chr19 | 3,947,739 | C | T | RR | RA | RR | RR | promoter | PtStettler14.19G030500 |
| Chr19 | 6,305,204 | C | T | RR | RR | RR | RA | intergenic | |
| Chr19 | 11,967,431 | C | T | RR | RR | RR | RA | TE | SINE2-1_PTr |
| Chr19 | 14,356,939 | A | T | RA | RR | RR | RR | mRNA | PtStettler14.19G100700 |
| Chr19 | 14,661,691 | A | T | RR | RA | RR | RR | mRNA | PtStettler14.19G103800 |

**Table S3.** Nucleotide mutation rate estimates for five filtering depths and multiple replicates. GS is the effective genome size.

| Depth | Replicate | Rate Estimate | Least square value | Tree 13 Effective GS | Tree 14 Effective GS |
|---|---|---|---|---|---|
| 20 | 1 | 1.33E-10 | 9.24E-13 | 40,388,007 | 54,998,919 |
| 20 | 2 | 1.17E-10 | 7.46E-13 | 40,388,007 | 54,998,919 |
| 20 | 3 | 1.22E-10 | 9.21E-13 | 40,388,007 | 54,998,919 |
| 25 | 1 | 1.04E-10 | 1.01E-12 | 44,900,782 | 58,223,515 |
| 25 | 2 | 1.10E-10 | 8.72E-13 | 44,900,782 | 58,223,515 |
| 25 | 3 | 1.17E-10 | 1.10E-12 | 44,900,782 | 58,223,515 |
| 30 | 1 | 1.07E-10 | 9.88E-13 | 38,229,529 | 49,555,495 |
| 30 | 2 | 1.14E-10 | 9.26E-13 | 38,229,529 | 49,555,495 |
| 30 | 3 | 1.12E-10 | 9.54E-13 | 38,229,529 | 49,555,495 |
| 35 | 1 | 9.52E-11 | 1.10E-12 | 32,546,285 | 40,561,041 |
| 35 | 2 | 9.44E-11 | 1.17E-12 | 32,546,285 | 40,561,041 |
| 35 | 3 | 9.34E-11 | 1.21E-12 | 32,546,285 | 40,561,041 |
| 40 | 1 | 9.03E-11 | 1.07E-12 | 24,989,388 | 28,972,798 |
| 40 | 2 | 8.94E-11 | 1.08E-12 | 24,989,388 | 28,972,798 |
| 40 | 3 | 8.95E-11 | 8.26E-13 | 24,989,388 | 28,972,798 |
| 45 | 1 | 7.03E-11 | 9.14E-13 | 18,264,725 | 19,351,957 |
| 45 | 2 | 7.44E-11 | 8.85E-13 | 18,264,725 | 19,351,957 |
| 45 | 3 | 8.15E-11 | 8.75E-13 | 18,264,725 | 19,351,957 |

**Table S4.** Total PacBio sequencing output for the branches used for structural variation analysis.

| Tree | Branch Name | Library Code | Total Reads | Total Basepairs | Single Pass Reads | Single Pass Basepairs | 20kb Reads | 20kb Basepairs |
|---|---|---|---|---|---|---|---|---|
| Tree 13 | 13.1 | PBAU | 3,649,818 | 39,102,995,683 | 2,980,747 | 35,239,027,694 | 632,022 | 17,945,029,823 |
| Tree 13 | 13.2 | PBAW | 1,457,219 | 15,778,516,050 | 1,013,763 | 13,102,813,747 | 279,681 | 7,600,620,682 |
| Tree 13 | 13.3 | PBAT | 3,863,624 | 39,855,580,888 | 3,076,499 | 35,493,092,219 | 624,581 | 17,880,248,612 |
| Tree 13 | 13.5 | PAZF | 2,477,499 | 20,286,559,665 | 2,185,822 | 18,741,157,420 | 288,846 | 8,101,551,515 |
| Tree 14 | 14.2 | PAXN | 1,934,882 | 20,857,551,670 | 1,665,573 | 19,342,446,966 | 344,342 | 10,224,080,227 |
| Tree 14 | 14.3 | PAXL | 3,838,857 | 32,981,037,681 | 3,195,080 | 29,347,720,852 | 396,508 | 10,688,170,010 |
| Tree 14 | 14.4 | PAYK | 3,624,069 | 28,300,147,045 | 3,142,139 | 26,054,497,423 | 308,645 | 8,204,269,654 |
| Tree 14 | 14.5 | PAZH | 3,549,237 | 28,894,702,695 | 3,115,179 | 26,761,563,517 | 340,884 | 9,183,104,989 |
| | | mean value | 3,049,401 | 28,257,136,422 | 2,546,850 | 25,510,289,980 | 401,939 | 11,228,384,439 |
| | | total value | 24,395,205 | 226,057,091,377 | 20,374,802 | 204,082,319,838 | 3,215,509 | 89,827,075,512 |

**Table S5.** Counts of SVs separated by type and size. Count is the mean of four replicates of 'pbsv call' with standard deviation in parentheses.

| | > 20bp | > 100bp | > 1kb | > 5kb | > 10kb | > 25kb | > 50kb |
|---|---|---|---|---|---|---|---|
| Deletion | 10,466.25 (26.42) | 4,539.75 (19.43) | 841.25 (8.73) | 208.0 (5.10) | 83.25 (1.71) | 25.75 (1.26) | 11 (0) |
| Insertion | 6,702.25 (39.58) | 3,920.75 (8.77) | 617.75 (3.86) | 28.25 (0.5) | 0 (0) | 0 (0) | 0 (0) |
| Duplication | 645 (6.58) | 297 (1.15) | 88.5 (0.58) | 59 (0.82) | 39 (0) | 14 (0) | 5 (0) |
| Inversion | 3 (0) | 3 (0) | 1 (0) | 0 (0) | 0 (0) | 0 (0) | 0 (0) |

**Table S6.** Support for SV designation of random subset of SVs based on visual evaluation of read mapping patterns in IGV. Percentages, by row, in parentheses.

| | | Total | Read Mapping Support | | |
|---|---|---|---|---|---|
| | | | Strong | Moderate | Weak |
| By type | All tested SVs | 85 | 59 (0.694) | 14 (0.165) | 12 (0.141) |
| | deletions (DEL) | 41 | 30 (0.732) | 8 (0.195) | 3 (0.073) |
| | duplications (DUP) | 34 | 19 (0.559) | 6 (0.176) | 0 (0.0) |
| | insertions (INS) | 20 | 20 (1.0) | 0 (0.0) | 0 (0.0) |
| By size | >50kb | 15 | 6 (0.400) | 7 (0.467) | 2 (0.133) |
| | 10kb-50kb | 20 | 8 (0.400) | 4 (0.200) | 8 (0.400) |
| | 1kb-10kb | 30 | 27 (0.900) | 1 (0.033) | 2 (0.067) |
| | 20bp-1kb | 30 | 28 (0.933) | 2 (0.067) | 0 (0.0) |
| type & size | DEL > 50kb | 11 | 5 (0.455) | 5 (0.455) | 1 (0.091) |
| | DEL 10kb-50kb | 10 | 5 (0.500) | 3 (0.300) | 2 (0.200) |
| | DEL 1kb-10kb | 10 | 10 (1.0) | 0 (0.0) | 0 (0.0) |
| | DEL 20bp-1kb | 10 | 10 (1.0) | 0 (0.0) | 0 (0.0) |
| | DUP >50kb | 4 | 1 (0.250) | 2 (0.500) | 1 (0.250) |
| | DUP 10kb-50kb | 10 | 1 (0.100) | 3 (0.300) | 6 (0.600) |
| | DUP 1kb-10kb | 10 | 1 (0.100) | 7 (0.700) | 2 (0.200) |
| | DUP 20bp-1kb | 10 | 8 (0.800) | 2 (0.200) | 0 (0.0) |
| | INS 1kb-10kb | 10 | 10 (1.0) | 0 (0.0) | 0 (0.0) |
| | INS 20bp-1kb | 10 | 10 (1.0) | 0 (0.0) | 0 (0.0) |

**Table S7.** Whole-genome bisulfite sequencing summary statistics.

| Branch | Input reads | Non-conversion (%) | Uniquely mapped reads | Percent mapped reads | Genome coverage |
|---|---|---|---|---|---|
| 13.1 | 195,656,607 | 0.152 | 105,029,741 | 53.68 | 40.16 |
| 13.2 | 196,639,972 | 0.124 | 106,408,336 | 54.11 | 40.69 |
| 13.3 | 208,799,824 | 0.127 | 113,022,558 | 54.13 | 43.22 |
| 13.5 | 183,361,979 | 0.124 | 99,746,602 | 54.40 | 38.14 |
| 14.2 | 199,024,702 | 0.158 | 107,727,193 | 54.13 | 41.20 |
| 14.3 | 187,580,466 | 0.146 | 102,282,378 | 54.53 | 39.11 |
| 14.4 | 204,539,730 | 0.155 | 111,007,228 | 54.27 | 42.45 |
| 14.5 | 213,186,674 | 0.149 | 114,199,385 | 53.57 | 43.67 |

**Table S8.** Calculated epimutation rates by sequence context and genomic feature. Alpha is the rate of gaining methylation. Beta is the rate of losing methylation. F-test compares the neutral model (degrees of freedom 23) vs null model (d.o.f. 27).

| Sequence | Feature | Alpha | Beta | Beta / alpha | F-test | P-value |
|---|---|---|---|---|---|---|
| CG | single | 1.70E-06 | 5.80E-06 | 3.40 | 154.20 | 2.97E-16 |
| CG | regions | 2.10E-06 | 6.10E-06 | 2.80 | 78.60 | 4.45E-13 |
| CG | mRNA | 2.50E-06 | 2.10E-05 | 8.40 | 298.20 | 1.88E-19 |
| CG | intergenic | 1.60E-06 | 5.60E-06 | 3.50 | 163.00 | 1.6E-16 |
| CG | promoter | 1.10E-06 | 8.00E-06 | 7.30 | 62.50 | 5.1E-12 |
| CG | TE | 7.50E-07 | 2.80E-07 | 0.37 | 28.10 | 1.46E-08 |
| CHG | single | 3.35E-07 | 4.07E-06 | 12.14 | 33.47 | 2.78E-09 |
| CHG | regions | 7.00E-07 | 3.90E-06 | 5.60 | 20.90 | 2.17E-07 |
| CHG | mRNA | 2.59E-07 | 3.82E-05 | 147.66 | 17.83 | 8.70E-07 |
| CHG | intergenic | 3.65E-07 | 4.06E-06 | 11.12 | 39.18 | 5.95E-10 |
| CHG | promoter | 3.59E-07 | 6.61E-06 | 18.44 | 29.89 | 8.25E-09 |
| CHG | TE | 3.23E-07 | 2.71E-07 | 0.84 | 9.67 | 9.73E-05 |
| CHH | single | 1.50E-08 | 3.60E-06 | 244.60 | 25.20 | 3.98E-08 |

| | | | | | | |
|---|---|---|---|---|---|---|
| CHH | regions | 4.80E-08 | 6.80E-06 | 141.70 | 5.17 | 4.00E-03 |
| CHH | mRNA | - | - | - | 0.005 | 0.99 |
| CHH | intergenic | - | - | - | 0.034 | 0.99 |
| CHH | promoter | - | - | - | 0.086 | 0.98 |
| CHH | TE | - | - | - | 0.219 | 0.99 |

**Table S9.** mRNA-seq library and mapping statistics.

| Branch | Input basepairs | Input reads | Reads map uniquely | Reads map >1 time | Total reads mapped | Overall mapping rate |
|---|---|---|---|---|---|---|
| 13.1 r1 | 23,869,710 | 47,739,420 | 40,907,228 | 5,529,951 | 46,437,179 | 97.39 |
| 13.1 r2 | 27,985,450 | 55,970,900 | 49,284,675 | 4,995,428 | 54,280,103 | 97.09 |
| 13.1 r3 | 29,318,523 | 58,637,046 | 45,811,779 | 11,277,278 | 57,089,057 | 97.47 |
| 13.2 r1 | 28,007,596 | 56,015,192 | 48,747,233 | 5,643,861 | 54,391,094 | 97.22 |
| 13.2 r2 | 26,738,198 | 53,476,396 | 46,259,530 | 5,615,821 | 51,875,351 | 97.12 |
| 13.2 r3 | 25,412,356 | 50,824,712 | 39,664,512 | 9,738,575 | 49,403,087 | 97.30 |
| 13.3 r1 | 28,057,987 | 56,115,974 | 51,405,968 | 3,022,313 | 54,428,281 | 97.10 |
| 13.3 r2 | 28,194,300 | 56,388,600 | 41,114,607 | 13,778,165 | 54,892,772 | 97.44 |
| 13.3 r3 | 24,355,844 | 48,711,688 | 38,376,276 | 8,888,828 | 47,265,104 | 97.13 |
| 13.5 r1 | 23,303,580 | 46,607,160 | 41,855,341 | 3,313,629 | 45,168,970 | 97.04 |
| 13.5 r2 | 22,709,851 | 45,419,702 | 40,553,261 | 3,636,067 | 44,189,328 | 97.41 |
| 13.5 r3 | 23,718,622 | 47,437,244 | 40,780,564 | 5,355,794 | 46,136,358 | 97.36 |
| 14.2 r1 | 28,117,599 | 56,235,198 | 50,070,189 | 4,261,492 | 54,331,681 | 96.68 |
| 14.2 r2 | 28,597,331 | 57,194,662 | 50,449,544 | 4,607,830 | 55,057,374 | 96.35 |
| 14.2 r3 | 28,579,533 | 57,159,066 | 45,029,115 | 10,145,808 | 55,174,923 | 96.59 |
| 14.3 r1 | 28,560,700 | 57,121,400 | 48,122,274 | 6,169,444 | 54,291,718 | 95.11 |
| 14.3 r2 | 29,843,996 | 59,687,992 | 35,935,530 | 20,880,835 | 56,816,365 | 95.26 |
| 14.3 r3 | 29,637,111 | 59,274,222 | 51,173,261 | 4,854,351 | 56,027,612 | 94.60 |
| 14.4 r1 | 26,168,420 | 52,336,840 | 42,802,511 | 7,078,556 | 49,881,067 | 95.44 |

| Branch | Input basepairs | Input reads | Reads map uniquely | Reads map >1 time | Total reads mapped | Overall mapping rate |
|---|---|---|---|---|---|---|
| 14.4 r2 | 29,913,292 | 59,826,584 | 45,546,254 | 12,513,281 | 58,059,535 | 97.08 |
| 14.4 r3 | 31,980,446 | 63,960,892 | 52,468,827 | 9,845,842 | 62,314,669 | 97.46 |
| 14.5 r1 | 31,008,774 | 62,017,548 | 50,349,762 | 9,728,826 | 60,078,588 | 96.92 |
| 14.5 r2 | 33,774,842 | 67,549,684 | 58,710,885 | 6,931,613 | 65,642,498 | 97.22 |
| 14.5 r3 | 28,940,195 | 57,880,390 | 47,595,291 | 8,755,418 | 56,350,709 | 97.40 |

**Table S10.** Genomic libraries included in the *Populus trichocarpa* var. *Stettler14* genome assembly and their respective assembled sequence coverage levels in the final release. *Average read length of PacBio reads.

| Sequencing platform | Average Read/Insert Size | Read number | Assembled sequence coverage (x) |
|---|---|---|---|
| Illumina | 400 +- 50 | 906,280,916 | 349 |
| PacBio | 10,477* | 4,887,084 | 118.58 |

**Table S11.** PacBio library statistics for total yield of the 64 chips included in the *Populus trichocarpa* var. *Stettler14* genome assembly and their respective assembled sequence coverage levels.

| Cutoff | Number of Reads | Basepairs | Average Read Length | Coverage |
|---|---|---|---|---|
| 0 | 4,887,084 | 59,290,798,933 | 10,477 | 118.58x |
| 1,000 | 4,682,541 | 59,175,547,228 | 10,967 | 118.35x |
| 2,000 | 4,427,480 | 58,791,470,782 | 11,595 | 117.58x |
| 3,000 | 4,164,497 | 58,134,477,994 | 12,263 | 116.27x |
| 4,000 | 3,910,519 | 57,246,542,875 | 12,936 | 114.49x |
| 5,000 | 3,668,457 | 56,158,151,033 | 13,604 | 112.32x |
| 6,000 | 3,435,698 | 54,878,877,846 | 14,273 | 109.76x |
| 7,000 | 3,208,490 | 53,402,831,014 | 14,964 | 106.81x |
| 8,000 | 2,985,015 | 51,727,059,646 | 15,673 | 103.45x |
| 9,000 | 2,763,432 | 49,843,715,135 | 16,415 | 99.69x |
| 10,000 | 2,545,360 | 47,772,740,258 | 17,183 | 95.55x |

| Cutoff | Number of Reads | Basepairs | Average Read Length | Coverage |
|---|---|---|---|---|
| 11,000 | 2,334,628 | 45,560,908,179 | 17,961 | 91.12x |
| 12,000 | 2,133,544 | 43,249,315,386 | 18,737 | 86.50x |
| 13,000 | 1,943,610 | 40,876,304,352 | 19,508 | 81.75x |
| 14,000 | 1,764,446 | 38,458,530,384 | 20,265 | 76.92x |
| 15,000 | 1,598,408 | 36,052,017,988 | 21,006 | 72.10x |
| 16,000 | 1,443,006 | 33,644,348,692 | 21,737 | 67.29x |
| 17,000 | 1,298,206 | 31,255,941,594 | 22,456 | 62.51x |
| 18,000 | 1,162,260 | 28,877,602,421 | 23,186 | 57.76x |
| 19,000 | 1,033,940 | 26,504,349,161 | 23,938 | 53.01x |

**Table S12.** Summary statistics of the raw output of the MECAT whole genome shotgun assembly. The table shows total contigs and total assembled base pairs for each set of scaffolds greater than the size listed in the left-most column.

| Minimum scaffold length | Number of scaffolds | Number of contigs | Scaffold Size | Basepairs | % Non-gap Basepairs |
|---|---|---|---|---|---|
| 5 Mb | 28 | 28 | 245,582,290 | 245,582,290 | 100 |
| 2.5 Mb | 49 | 49 | 322,995,609 | 322,995,609 | 100 |
| 1 Mb | 77 | 77 | 371,688,077 | 371,688,077 | 100 |
| 500 Kb | 103 | 103 | 388,187,453 | 388,187,453 | 100 |
| 250 Kb | 188 | 188 | 415,811,552 | 415,811,552 | 100 |
| 100 Kb | 955 | 955 | 523,394,177 | 523,394,177 | 100 |
| 50 Kb | 2,985 | 2,985 | 664,242,744 | 664,242,744 | 100 |
| 25 Kb | 3,665 | 3,665 | 693,169,682 | 693,169,682 | 100 |
| 10 Kb | 3,693 | 3,693 | 693,764,729 | 693,764,729 | 100 |
| 5 Kb | 3,693 | 3,693 | 693,764,729 | 693,764,729 | 100 |
| 2.5 Kb | 3,693 | 3,693 | 693,764,729 | 693,764,729 | 100 |
| 1 Kb | 3,693 | 3,693 | 693,764,729 | 693,764,729 | 100 |
| 0 bp | 3,693 | 3,693 | 693,764,729 | 693,764,729 | 100 |

**Table S13.** Divergence ages between branches used to compute mutation and epimutation rates. Time of sample 1 and 2 are the number of years of growth since the tree began. Time of ancestor is the number of years of growth to the samples most recent common branch point.

| Sample 1 | Sample 2 | Time of sample 1 | Time of sample 2 | Time most common ancestor | Delta t | D.value |
|---|---|---|---|---|---|---|
| 13.1 | 13.2 | 328 | 327 | 286 | 83 | 0.00235717 |
| 13.1 | 13.3 | 328 | 328 | 258 | 140 | 0.00295603 |
| 13.1 | 13.5 | 328 | 297 | 217 | 191 | 0.00283421 |
| 13.1 | 14.2 | 328 | 324 | 0 | 652 | 0.00428722 |
| 13.1 | 14.3 | 328 | 324 | 0 | 652 | 0.00416599 |
| 13.1 | 14.4 | 328 | 287 | 0 | 615 | 0.00432367 |
| 13.1 | 14.5 | 328 | 287 | 0 | 615 | 0.00404655 |
| 13.2 | 13.3 | 327 | 328 | 258 | 139 | 0.00284635 |
| 13.2 | 13.5 | 327 | 297 | 217 | 190 | 0.00269338 |
| 13.2 | 14.2 | 327 | 324 | 0 | 651 | 0.00426179 |
| 13.2 | 14.3 | 327 | 324 | 0 | 651 | 0.00412762 |
| 13.2 | 14.4 | 327 | 287 | 0 | 614 | 0.00421219 |
| 13.2 | 14.5 | 327 | 287 | 0 | 614 | 0.00399516 |
| 13.3 | 13.5 | 328 | 297 | 217 | 191 | 0.00277805 |
| 13.3 | 14.2 | 328 | 324 | 0 | 652 | 0.0043418 |
| 13.3 | 14.3 | 328 | 324 | 0 | 652 | 0.00417754 |
| 13.3 | 14.4 | 328 | 287 | 0 | 615 | 0.00425596 |
| 13.3 | 14.5 | 328 | 287 | 0 | 615 | 0.00404867 |
| 13.5 | 14.2 | 297 | 324 | 0 | 621 | 0.00416148 |
| 13.5 | 14.3 | 297 | 324 | 0 | 621 | 0.00397476 |
| 13.5 | 14.4 | 297 | 287 | 0 | 584 | 0.00404067 |
| 13.5 | 14.5 | 297 | 287 | 0 | 584 | 0.00379661 |
| 14.2 | 14.3 | 324 | 324 | 283 | 82 | 0.00234308 |

| Sample 1 | Sample 2 | Time of sample 1 | Time of sample 2 | Time most common ancestor | Delta t | D.value |
|---|---|---|---|---|---|---|
| 14.2 | 14.4 | 324 | 287 | 247 | 117 | 0.00274387 |
| 14.2 | 14.5 | 324 | 287 | 215 | 181 | 0.00273497 |
| 14.3 | 14.4 | 324 | 287 | 247 | 117 | 0.00248307 |
| 14.3 | 14.5 | 324 | 287 | 215 | 181 | 0.00248201 |
| 14.4 | 14.5 | 287 | 287 | 215 | 144 | 0.00253796 |

### mutSOMA: Estimating somatic mutation rates from high-throughput sequencing data in trees

We developed *mutSOMA*, a computational method for estimating somatic mutation rates from high-throughput sequencing data in trees. The method treats the tree branching structure as an intra-organism phylogeny of somatic lineages. Its analytical framework builds on ideas introduced in van der Graaf et al. (2015) and Shahryary et al. 2019 (co-submission). Software implementing the method can be found at (https://github.com/jlab-code/mutSOMA).

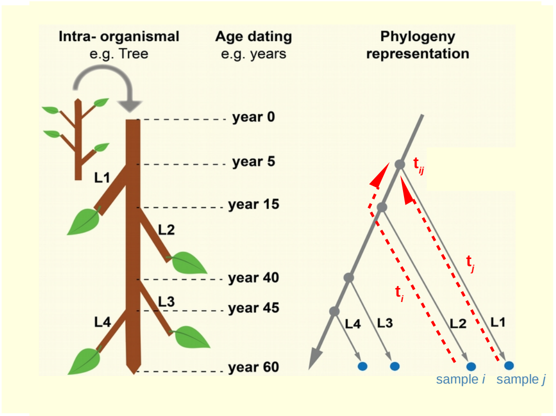


**Figure 1**: Long-lived perennials, such as trees, can be viewed as a natural mutation accumulation system. In this case, the tree branching structure can be treated as an intra-organismal phylogeny of somatic lineages that carry information about the mutational history of each branch. Re-sequencing data is obtained from leaf samples of selected branches. *mutSOMA* uses the genotype data of the samples along with the coring data to estimate the per year rate of somatic mutations. L1, L2, L3, L4 denote the branches of the tree; blue circles denote sequenced samples; grey circles denote branch points. Highlighted are samples *i* and *j* and the corresponding branch ages $t_i$, $t_j$ as well as the age of the most recent common branch point $t_{ij}$.

#### Calculating genetic divergence

We start from the variant calls (i.e. .vcf files) obtained from different branches of the tree (**Figure 1**). For the i-*th* sample ($i = 1, \ldots, M$) we let $g_{ik}$ be the observed genotype at the k-*th* locus ($k = 1, \ldots, N$), where $N$ is the effective genome size (i.e. the total number of bases with sufficient coverage). With four possible nucleotides (A, C, T, G), $g_{ik}$ can have 16 possible genotypes in a diploid genome, 4 homozygous (A|A, T|T, C|C, G|G) and 12 heterozygous (A|G, A|T, ..., G|C). The $\cdot|\cdot$ notation refers to the nucleotide on the forward (+) strand on each of the two homologous chromosomes. Using this coding, we calculate the genetic divergence, $D$, between any two samples $i$ and $j$ as follows:

$$D_{ij} = \sum_{k=1}^{N} I(g_{ik}, g_{jk}) N^{-1}, \tag{1}$$

where $I(\cdot)$ is an indicator function, such that, $I(\cdot) = 0$ if the two samples share no alleles at locus k (e.g. A|A and G|G), 0.5 if they share one (e.g. A|A and A|G), and 1 if they share both alleles (e.g. A|A and A|A). We suppose that $D_{ij}$ is related to the developmental divergence time of samples $i$ and $j$ through a somatic mutation model. The divergence times (in years) are calculated from the coring data (**Figure 1**).

#### Modelling age-dependent genetic divergence

We model the time-dependent genetic divergence between samples using

$$D_{ij} = c + D^{\bullet}_{ij}(M_{\Theta}) + \epsilon_{ij}. \tag{2}$$

Here $\epsilon_{ij} \sim N(0, \sigma^2)$ is the normally distributed residual error, $c$ is the intercept, and $D^{\bullet}_{ij}(M_{\Theta})$ is the expected genetic divergence between samples $i$ and $j$ as a function of an underlying mutation model $M(\cdot)$ with parameter vector $\Theta$. Parameter vector $\Theta$ contains the unknown mutation rate $\gamma$ and the unknown level of heterozygosity $\delta$ of the 'founder cells' of the tree (**see Figure 1**). The estimation of the residual variance in the model accounts for the fact that part of the observed genetic divergence between any two samples is driven by genotyping errors. We have that

$$\begin{aligned} D^{\bullet}_{ij}(M_{\Theta}) &= \sum_{n \in v} \sum_{l \in v} \sum_{m \in v} I(l, m) \\ &\cdot \; Pr(g_{ik} = l, g_{jk} = m | g_{ijk} = n, M_{\Theta}) \\ &\cdot \; Pr(g_{ijk} = n | M_{\Theta}), \end{aligned}$$

where $g_{ijk}$ is the genotype at the $k$ locus of the the most recent progenitor cells that are developmentally shared between samples $i$ and $j$, and $v \in \{\text{A|A, T|T, C|C, \ldots, G|T}\}$. Since the two samples are conditionally independent, we can further write:

$$Pr(g_{ik}, g_{jk}|g_{ijk}, M_\Theta) = Pr(g_{ik}|g_{ijk}, M_\Theta) \cdot Pr(g_{jk}|g_{ijk}, M_\Theta).$$

To be able to evaluate these conditional probabilities it is necessary to posit an explicit form for the somatic mutation model, $M_\Theta$. To motivate this, we define $\mathbf{G}_{(16\times16)}$ to be a $16 \times 16$ transition matrix, which summarizes the probability of transitioning from genotype $l$ to $m$ in the time interval $[t, t+1]$. $\mathbf{G}$ can be written in the following partitioned form:

$$\mathbf{G}_{(16\times16)} = \left(\begin{array}{c|c} \mathbf{T1}_{(4\times4)} & \mathbf{T2}_{(4\times12)} \\ \hline \mathbf{T3}_{(12\times4)} & \mathbf{T4}_{(12\times12)} \end{array}\right)$$

where sub-matrices **T1**, **T2**, **T3** and **T4** contain the transition probabilities between homozygous to homozygous, homozygous to heterozygous, heterozygous to homozygous and heterozygous to heterozygous genotypes, respectively. Explicit elements of each of these matrices can be worked out and hold for both somatic and clonally propagated systems. As there is no genetic segregation, the elements of this matrix are only governed by the mutation rate $\gamma$. For instance, symmetrical sub-matrix **T1** is

$$\mathbf{T1}_{(4\times4)} = \begin{array}{c} \begin{array}{cccc} \text{A|A (t+1)} & \text{C|C (t+1)} & \text{T|T (t+1)} & \text{G|G (t+1)} \end{array} \\ \begin{bmatrix} (1-\gamma)^2 & \frac{1}{9}\gamma^2 & \frac{1}{9}\gamma^2 & \frac{1}{9}\gamma^2 \\ \cdot & (1-\gamma)^2 & \frac{1}{9}\gamma^2 & \frac{1}{9}\gamma^2 \\ \cdot & \cdot & (1-\gamma)^2 & \frac{1}{9}\gamma^2 \\ \cdot & \cdot & \cdot & (1-\gamma)^2 \end{bmatrix} \end{array} \begin{array}{l} \text{A|A (t)} \\ \text{C|C (t)} \\ \text{T|T (t)} \\ \text{G|G (t)} \end{array}$$

, and **T4** is

$$\mathbf{T4}_{(12\times 12)} = \begin{array}{c} \begin{array}{cccc} \text{A|C (t+1)} & \text{A|T (t+1)} & \cdot & \text{G|T (t+1)} \end{array} \\ \left[\begin{array}{cccc} (1-\gamma)^2 & \frac{1}{3}(1-\gamma)\gamma & \cdot & \frac{1}{9}\gamma^2 \\ \cdot & \cdot & \cdot & \cdot \\ \cdot & \cdot & \cdot & \frac{1}{9}\gamma^2 \\ \cdot & \cdot & \cdot & (1-\gamma)^2 \end{array}\right] \end{array} \begin{array}{l} \\ \text{A|C (t)} \\ \text{A|T (t)} \\ \cdot \\ \text{G|T (t)} \end{array}$$

.

Based on Markov chain theory, the conditional probability $Pr(g_{ik}|g_{ijk}, M_\Theta)$ can then be expressed in terms of **G** as follows:

$$\sum_{n\in v} Pr(g_{ik} = v_r|g_{ijk} = n, M_\Theta) = \sum_{s=1}^{16} \left[\mathbf{G}^{t_i - t_{ij}}\right]_{rs}$$

where $r = 1, \ldots, 16$ is a fixed index corresponding to genotype vector {A|A, C|C, ..., G|T }, $t_i$ is the age of sample $i$ and $t_{ij}$ is the age of the most recent common branch point of samples $i$ and $j$, $(t_{ij} \leq t_i, t_j)$. Expressions for $Pr(g_{jk}|g_{ijk}, M_\Theta, t_j)$ can be derived accordingly, by simply replacing $t_i$ by $t_j$ in the above equation. Note that the calculation of these conditional probabilities requires repeated matrix multiplication. However, a direct evaluation of these equations is also possible using the fact that

$$\mathbf{G}^{t_i - t_{ij}} = \mathbf{p}\mathbf{V}^{t_i - t_{ij}}\mathbf{p}^{-1} \text{ and } \mathbf{G}^{t_j - t_{ij}} = \mathbf{p}\mathbf{V}^{t_j - t_{ij}}\mathbf{p}^{-1},$$

where **p** is the eigenvector of matrix **G** and **V** is a diagonal matrix of eigenvalues. Finally, to derive $D^{\bullet}_{ij}(M_\Theta)$, we also need to supply $Pr(g_{ijk} = n|M_\Theta)$; that is, the probability that any given locus $k$ in most recent shared progenitor cells of $i$ and $j$ is in state $n$ ($n \in$ {A|A, C|C, ..., G|T }). To do this, consider the genome of the hypothetical founder cell of tree at time $t = 1$, and let $\pi = [p_1\ p_2\ p_3\ \ldots\ p_{16}]$ be a row vector of probabilities corresponding to 16 possible genotypes, respectively. Using

Markov Chain theory we have

$$Pr(g_{ijk} = v_r | M_\Theta) \quad = \quad \left[\pi \, \mathbf{G}^{(t_{ij}-1)}\right]_r .$$

Assuming that *Populus trichocarpa* genome is at an evolutionary mutation equilibrium, we can obtained the probability elements of vector $\pi$ as follows

$$p_1 = \frac{x(A|A)}{N}(1-\delta), \qquad p_2 = \frac{x(C|C)}{N}(1-\delta), \qquad \ldots, \qquad p_{16} = \frac{x(G|T)}{12N}\delta$$

where $\delta \in [0,1]$ is the overall level of heterozygosity in the genome, $x(\cdot)$ is the frequency count of the loci with that particular genotype, and $N$ is the effective genome size.

#### Model inference

To obtain estimates for $\Theta$, we seek to minimize

$$\nabla \textstyle\sum_{q=1}^{M} \left(D_q - D_q^{\bullet}(M_\Theta) - c\right)^2 = \mathbf{0}, \tag{3}$$

where the summation is over all $M$ unique pairs of sequenced samples in the pedigree. Minimization is performed using the "Nelder-Mead" algorithm as part of the optimx package in R.

##### Confidence intervals

We obtain confidence intervals for the estimated model parameters by boostrapping the model residuals. The procedure has the following steps: 1. For the $q$th sample pair $q$ ($q = 1, \cdots, M$) we define a new response variable $B_q = \hat{D}_q + \hat{\epsilon}_k$, where $\hat{D}_q$ is the fitted divergence for the $q$th pair, and $\hat{\epsilon}_k$ is drawn at random and with replacement from the $1 \times M$ vector of fitted model residuals; 2. Refit the model using the new response variable, and obtain estimates for the model parameters. 3. Repeat steps 1. to 2. a large number of times to obtain a boostrap distribution. 4. Use the bootrap distribution from 3. to obtain empirical confidence intervals.
